# Functional connectivity and information pathways in the human entorhinal-hippocampal circuitry

**DOI:** 10.1101/2021.12.17.473123

**Authors:** Xenia Grande, Magdalena Sauvage, Andreas Becke, Emrah Düzel, David Berron

## Abstract

Cortical processing streams for item and contextual information come together in the entorhinal-hippocampal circuitry. Various evidence suggest that information-specific pathways organize the cortical – entorhinal interaction and the circuitry’s inner communication along the transversal axis. Here, we leveraged ultra-high field functional imaging and advance Maass, Berron et al. (2015) who report two functional routes segregating the entorhinal cortex (EC) and subiculum. Our data show specific scene processing in the functionally connected posterior-medial EC and distal subiculum. The regions of another route, that connects the anterior-lateral EC and a newly identified retrosplenial-based anterior-medial EC subregion with the CA1/subiculum border, process object and scene information similarly. Our results support topographical information flow in human entorhinal-hippocampal subregions with organized convergence of cortical processing streams and a unique route for contextual information. They characterize the functional organization of the circuitry and underpin its central role in memory function and pathological decline.

## Introduction

Entorhinal and hippocampal subregions form a critical functional circuitry that pulls cortical information together into cohesive representations (Eichenbaum et al., 2007; Ritchey et al., 2015). The interaction between the entorhinal-hippocampal circuitry and cortical information streams and the circuitry’s inner communication are key to the formation of these cohesive representations. Here we advance insight into how the human entorhinal cortex (EC) receives information from cortical streams and how information continues to flow from the EC towards the transversal hippocampal axis. These insights are relevant to our understanding of the circuitry’s fundamental role in cognitive functions such as episodic memory.

Two large-scale cortical information streams map onto different subdivisions of the EC. The two streams, that originate in the visual ‘What’ and ‘Where’ pathways, preferentially process item and contextual information in anterior-temporal cortical regions (i.e. orbitofrontal cortices, amygdalae and perirhinal cortices) versus posterior-medial cortical regions (i.e. retrosplenial and parahippocampal cortices), respectively (David Berron et al., 2018; Haxby et al., 1991; Ranganath & Ritchey, 2012; Ritchey et al., 2015; Ungerleider & Haxby, 1994). The way the EC communicates with different regions of these two cortical streams implies topographical differences in information processing within the EC. The medial EC subdivision preferentially communicates with the parahippocampal cortex. This parahippocampal source of cortical information is part of the contextual stream and in accordance, the medial EC seems to be tuned to process spatial or contextual information. The lateral EC preferentially communicates with the perirhinal cortex. The perirhinal cortex is a source of information from the item stream and in accordance, the lateral EC seems to process non-spatial or item information. These insights are derived from diverse studies on the anatomical wiring in rodents, functional connectivity in humans and functional studies in both species investigating various aspects of item versus contextual information processing (Neunuebel et al., 2013; Knierim et al., 2014; Reagh and Yassa, 2014; Zhang et al., 2014; Igarashi and Moser, 2015; Keene et al., 2016; Connor and Knierim, 2017; Berron et al., 2018). However, recently a more complex organization has been suggested. Unlike most conventional research that conceived the perirhinal and parahippocampal cortices as main information-specific inputs, recent anatomical findings show that the parahippocampal cortex communicates with the EC along its full extend, updating the conception of parallel perirhinal and parahippocampal input (Nilssen et al., 2019). These findings emphasize the retrosplenial cortex as a selective source to convey information from the posterior-medial stream towards the medial EC. This update fits recent insights suggesting an early convergence of spatial and non-spatial information in the rodent lateral EC (Doan et al., 2019). It evokes the question how these cortical sources of information uniquely map onto the EC and thereby contribute to the information pathways in the entorhinal-hippocampal circuitry.

Within the entorhinal-hippocampal circuitry, an important direct way of communication exists between the EC and hippocampal subiculum and CA1, which is characterized by two functional routes. A fully transversal organization of these routes is generally indicated in rodents where the anatomy and function show that spatial and non-spatial information is processed in two anatomically wired routes, the lateral EC – proximal subiculum – distal CA1 and the medial EC – distal subiculum – proximal CA1 route, respectively (Witter et al., 2017; note sparse functional evidence in the subiculum: Ku et al., 2017; Cembrowski et al., 2018; but frequent reports in the rodent CA1 region: Henriksen et al., 2010; Nakamura et al., 2013; Igarashi et al., 2014; Nakazawa et al., 2016; Beer et al., 2018). This aspect of inner entorhinal-hippocampal circuitry communication is less well studied in primates. For instance, the monkey anatomical wiring hints towards a longitudinal gradient on top of the transversal gradient with mainly anterior-lateral and posterior-lateral entorhinal portions projecting to the distal subiculum – proximal CA1 and proximal subiculum – distal CA1, respectively (Witter & Amaral, 2020), whereas in humans the functional connectivity pattern shows that the posterior-medial versus anterior-lateral entorhinal subregions preferentially connect to distal and proximal portions of the subiculum, respectively (Maass, Berron et al., 2015). Further functional evidence on the information flow in the human entorhinal-hippocampal circuitry is lacking. This calls for an investigation of the hypothesis from rodent research that information continues to flow in a segregated manner along the transversal human entorhinal-hippocampal axis.

To summarize, the cortical connectivity of the entorhinal-hippocampal circuitry and its related inner communication are still not fully understood. It is unclear first, in which subdivisions of the entorhinal-hippocampal circuitry item and context information are processed. Hence, the general connectivity patterns in the human entorhinal-hippocampal circuitry have not yet been directly related to information processing. Second, it is unclear where retrosplenial input maps onto the human EC as a source of the posterior-medial cortical system that processes contextual information. Therefor it is also unclear how that retrosplenial information is projected from the EC towards the hippocampus. Third, it remains elusive in how far a transversal functional segregation can be extended towards the human CA1 region in analogy to rodent literature. Note that a recent structural connectivity study is among the first studies in humans that examines retrosplenial input towards the EC (Syversen et al., 2021). This is to our knowledge also the first study that investigates the connectivity profile of the distal CA1 and thus suggests an expansion of the transversal connectivity pattern in the human brain as well. An unbiased assessment of information-specific functional connectivity and information-specific activity within the entorhinal and hippocampal subiculum and CA1 is, however, lacking.

How the entorhinal-hippocampal circuitry interacts with the cortex and conveys that information internally to process it, is also highly relevant from a clinical perspective as this circuitry is particularly vulnerable to tau pathology. Early cortical tau pathology seems to accumulate in Brodmann Area 35 or the perirhinal cortex, the lateral EC and can subsequently be found in the subiculum/CA1 border (David Berron et al., 2021; Lace et al., 2009), indicating a role of the connectivity with the proximal subiculum and distal CA1 for the anatomical progression of tau pathology. Indeed, an influential hypothesis suggests tau progression along functionally connected pathways in the human brain and associated memory decline (Adams et al., 2019; David Berron et al., 2020; Franzmeier et al., 2020; Maass et al., 2019; Vogel et al., 2020).

The central role of the entorhinal-hippocampal circuitry in memory function and the specific spread of Alzheimer’s disease pathology motivates us to advance insight into the information pathways within this circuitry and span existing knowledge from ex vivo animal research to in vivo functional imaging in humans. We leverage ultra-high resolution 7 Tesla functional imaging (fMRI) data and advance the earlier findings on human entorhinal subregions and the transversal intrinsic functional connectivity pattern in the subiculum (Maass, Berron et al., 2015). Here, we draw on entorhinal subregions that uniquely show functional connectivity with the retrosplenial, parahippocampal and perirhinal cortices. With respect to the perirhinal cortex, we also distinguish between Area 35 and Area 36 to allow for the translation of our findings to clinical research. In addition, we expand the functional investigation of entorhinal-hippocampal communication towards the human CA1 region to bring knowledge from animal research together with human research. Our unbiased functional connectivity approach likewise extends recent structural data (Syversen et al., 2021). Finally, we advance earlier findings by investigating functional connectivity together with associated biases for scene and object information processing in the same dataset (as aspects of contextual and item information processing, respectively). With these advances, we comprehensively examine how the entorhinal-hippocampal circuitry is embedded within large-scale cortical communication and how functionally and clinically relevant cortical sources contribute in a unique way to the information pathways within the entorhinal-hippocampal circuitry.

## Results

We seek to comprehensively investigate information pathways within the entorhinal-hippocampal circuitry and the contribution of cortical item and contextual processing streams. In an initial step, we identified where cortical sources map uniquely onto the entorhinal cortex (building upon Maass, Berron et al., 2015). The identified entorhinal subregions are based on their voxel’s preferred intrinsic functional connectivity with the retrosplenial cortex, parahippocampal cortex, perirhinal Area 36 or Area 35 regions (“sources”). Next, we evaluated the continuation of the functional connectivity streams within the entorhinal-hippocampal circuitry and examined the intrinsic functional connectivity pattern between the identified entorhinal subregions (“seeds”) and hippocampal subiculum and CA1 (again, building upon Maass, Berron et al., 2015). Therefore, temporal fluctuations of BOLD signal were correlated in a seed-to-voxel manner within each participant. The resulting statistical correlational maps were aligned between participants. Repeated measures ANOVAs were calculated on connectivity preferences with seeds and transversal segments as factors to determine statistical differences in connectivity topography.

Finally, we identified the corresponding bias in object (“item”) versus scene (“contextual”) information processing within the entorhinal subregions and along the hippocampal transversal axis, in the same dataset. Therefore, we extracted parameter estimates from a mnemonic discrimination task of object and scene information processing conditions from aligned statistical maps across participants. Repeated measures ANOVAs were calculated on parameter estimates in entorhinal subregions and transversal hippocampal segments to determine biases in information processing within the entorhinal-hippocampal circuitry.

In the following, we first describe the four obtained entorhinal seeds and display the intrinsic functional connectivity pattern with the entorhinal seed regions along the subiculum and CA1 transversal axes. Thereafter, we report the information processing characteristics of the entorhinal and hippocampal subregions. Note that all results have been obtained with independent analyses in the left and right hemispheres. The largely similar left hemisphere results can be found in supplementary I.

### Four cortical sources split the EC in anterior-lateral, posterior-medial, anterior-medial and posterior-lateral seeds

The four entorhinal subregions that we later used as seeds to determine the topography of entorhinal-hippocampal connectivity are based on intrinsic functional connectivity preferences with either the parahippocampal cortex, the retrosplenial cortex, Area 36 or Area 35. These cortical regions are in general concordance with Maass, Berron et al. (2015) but consider recent emphasis on the retrosplenial cortex as a source that conveys contextual information from the posterior-medial stream (Nilssen et al., 2019; Syversen et al., 2021) as well as potential clinical implications for anatomical Alzheimer’s tau pathology progression (Jacobs et al., 2018; Ziontz et al., 2021).

Based on functional connectivity preferences to the parahippocampal cortex, we obtained a largely posterior-medial entorhinal seed and based on preferences to the retrosplenial cortex a rather anterior-medial entorhinal seed. In contrast to these medially located entorhinal seeds we obtained lateral seeds based on connectivity preferences to the perirhinal Area 36 and Area 35. Anterior-lateral entorhinal voxels preferentially connected to Area 35 while posterior-lateral entorhinal voxels preferentially connected to Area 36. Note that both entorhinal seeds extended along the anterior to posterior axis such that the one based on connectivity with Area 35 progresses more along deep entorhinal portions (with a main focus anteriorly) and the one based on connectivity with Area 36 along superficial entorhinal portions (with a main focus posteriorly, see Figure 1).

**Figure 1.**
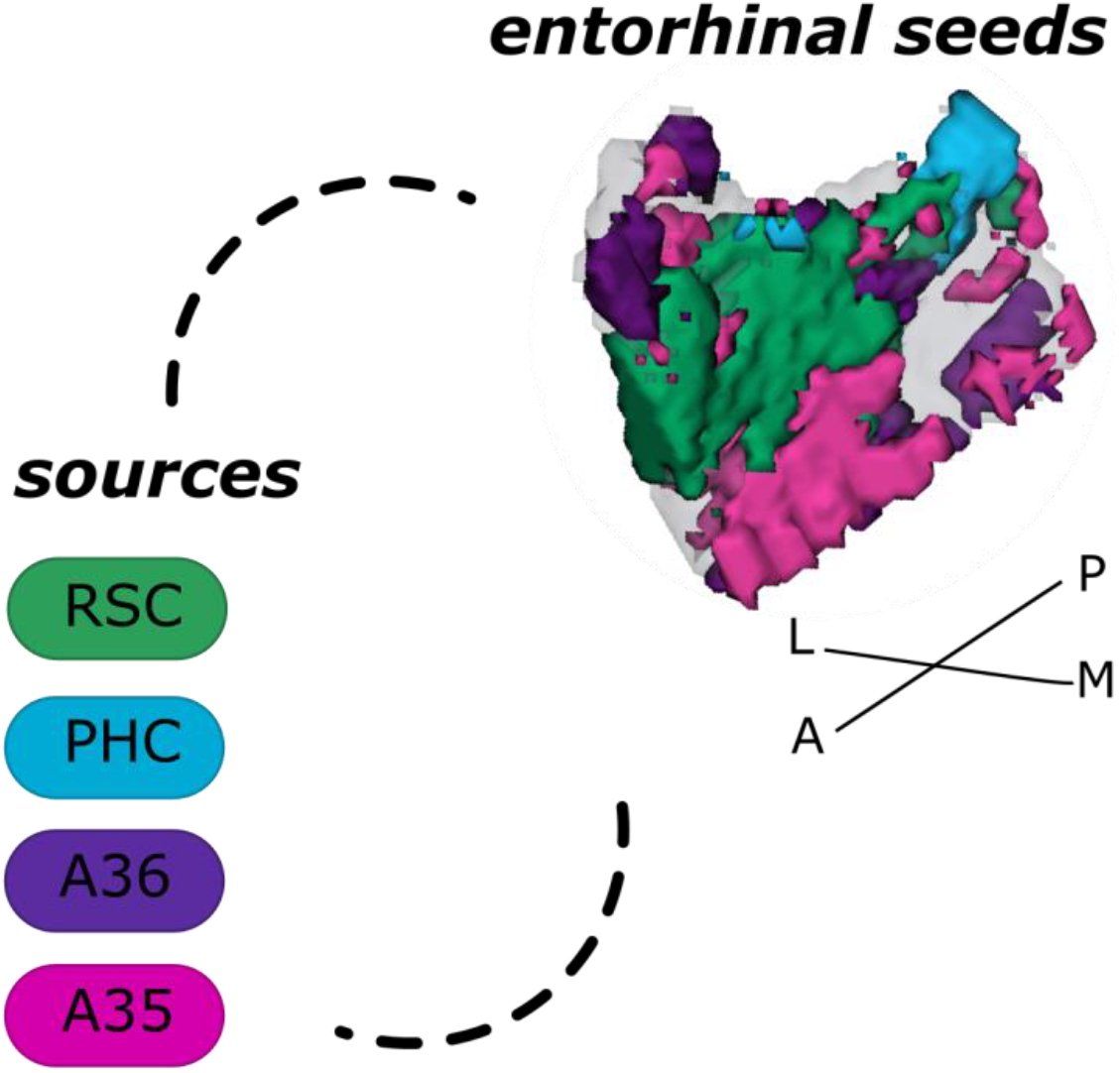
Entorhinal seed regions based on connectivity preferences to cortical regions. Displayed is the right entorhinal cortex as a 3D image with colored seed regions. The seed regions have been identified based on a source-to-voxel functional connectivity analysis and resulting connectivity preference to either the right retrosplenial (RSC, green) cortex, parahippocampal cortex (PHC, blue), Area 36 (A36, purple) or Area 35 (A35, pink) sources. Seed regions have been determined based on the thresholded (T > 3.1) maximum voxels across four one-sample t-tests at group level, one per source. M – medial; L – lateral; A – anterior; P – posterior.

### The distal subiculum functionally connects to the posterior-medial EC and the subiculum/CA1 border to anterior-medial and anterior-lateral EC

Following the characterization of entorhinal seeds, we focus on the functional connectivity between the entorhinal subregion and hippocampal subiculum and CA1 to advance the transversal functional connectivity profile described by Maass, Berron et al. (2015). When extracting estimates of connectivity preferences across individuals from proximal and distal hippocampal subfield segments for either entorhinal seed, repeated measures ANOVAs revealed significant seed X segments interaction effects along the transversal axis in the subiculum and in CA1 (see Figure 2; subiculum: F(12,372) = 19.561; p < .001; CA1: F(6,186) = 3.212; p = .024; FDR-corrected).

**Figure 2.**
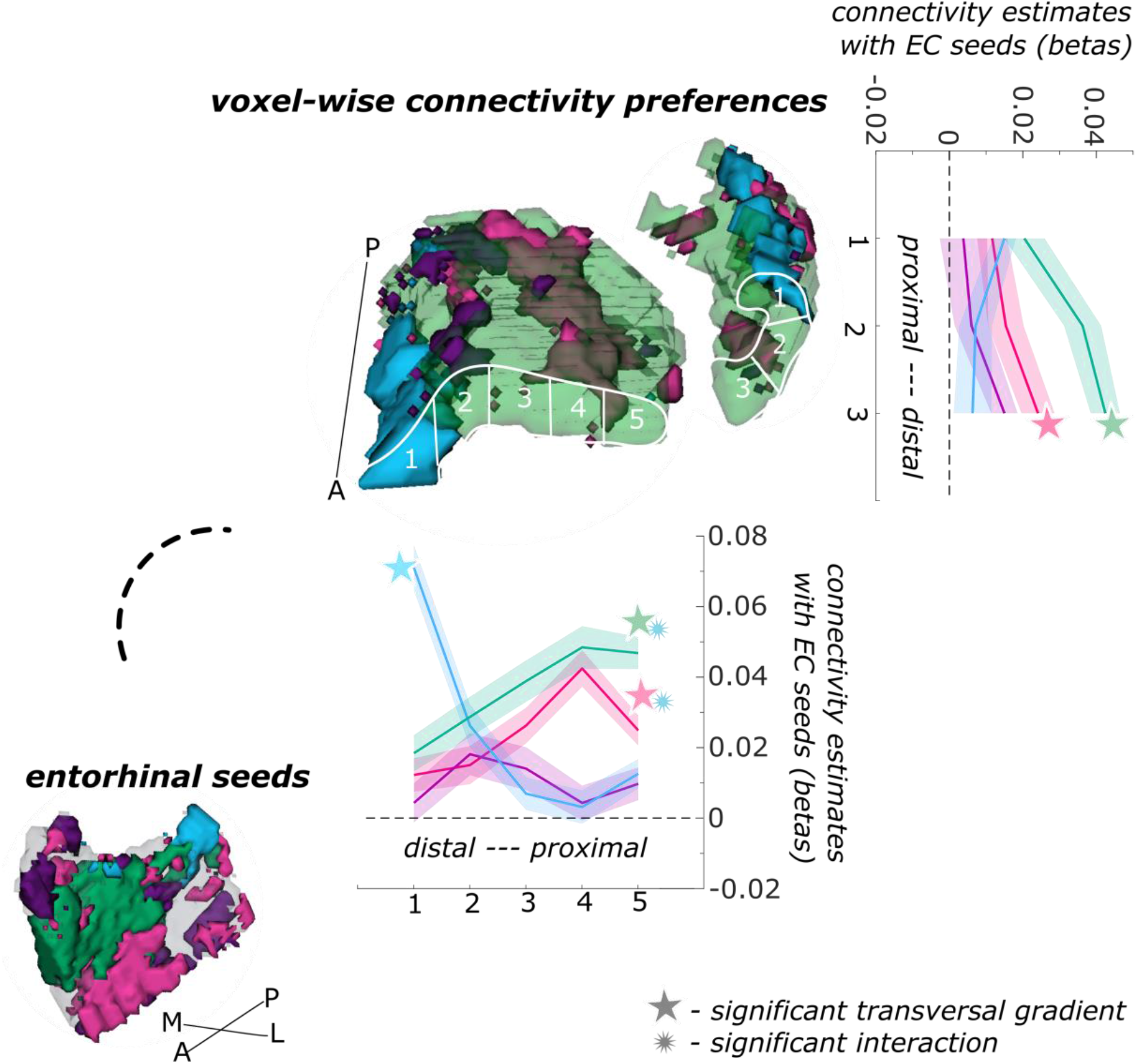
Functional connectivity preferences to entorhinal seeds along the subiculum and CA1 transversal axis. Displayed are the results of a seed-to-voxel functional connectivity analysis between the displayed right entorhinal seeds and the right subiculum and CA1 subregion. The 3D figure displays voxel-wise connectivity preferences to the entorhinal seeds (color coded to refer to the respective entorhinal seed) on group level. To display mean connectivity preferences across participants along the transversal axis, beta estimates were extracted and averaged from equally sized segments from proximal to distal ends (five segments in subiculum, three segments in CA1; schematized in white on the 3D figures) on each coronal slice and averaged along the longitudinal axis. Repeated measures ANOVAs revealed significant differences in connectivity estimates along the transversal axis in subiculum and CA1 with interaction effects in the subiculum. Displayed significances obtained by FDR-corrected post-hoc tests and refer to p < .05. Shaded areas in the graphs refer to standard errors of the mean. EC – entorhinal; M – medial; L – lateral; A – anterior; P – posterior.

In the subiculum, additional repeated measures ANOVAs showed that the anterior-lateral (F(4,124) = 8.913; p < .001), anterior-medial (F(4,124) = 10.538; p < .001) and posterior-medial (F(4,124) = 42.201; p < .001) entorhinal seeds displayed a significant main effect across the transversal subiculum segments. These differential functional connectivity preferences across the transversal axis interacted significantly in a subsequent repeated measures ANOVA (posterior-medial versus anterior-medial seed preference interaction: F(4,124) = 46.452; p < .001; posterior-medial versus anterior-lateral seed preference interaction: F(4,124) = 35.208; p <.001). This pattern provides statistical evidence for an increase in preferential functional connectivity with the posterior-medial entorhinal seed towards the distal portion of the subiculum while the preferential functional connectivity with the anterior-lateral as well as the anterior-medial entorhinal seeds rather increased towards the proximal portion of the subiculum.

In hippocampal CA1, additional repeated measures ANOVAs showed that the connectivity preference towards the anterior-medial entorhinal seed displays a significant main effect across the transversal CA1 axis (F(2,62) = 10.489; p < .001). In the distal CA1, the preferential functional connectivity with the posterior-medial entorhinal seed was higher than in the proximal portion of CA1. In the right CA1, a similar but weaker transversal pattern was observed for connectivity preferences with the anterior-lateral entorhinal cortex (F(2,62) = 4.146; p = .041; FDR-corrected; note in the left hemisphere a comparable transversal pattern was observed for the posterior- and anterior-medial portions of the entorhinal cortex, see supplementary I).

Thus, in the entorhinal-hippocampal circuitry, voxels in the distal subiculum preferentially connected functionally with the posterior-medial entorhinal portion whereas voxels in the subiculum/CA1 border preferentially connected with the anterior-medial as well as the anterior-lateral EC portion.

### The distal subiculum and posterior-medial EC specifically process object information while the other subregions process object and scene information similarly

Besides the intrinsic functional connectivity patterns within the entorhinal-hippocampal circuitry, we also examined the characteristics of object and scene information processing.

We first focus on the entorhinal seed regions. When extracting task-related parameter estimates for object and scene information processing, a repeated measures ANOVA showed a significant interaction between region and information type (object versus scene, F(3,93) = 20.9267; p < .001). Post-hoc t-tests revealed that only in the posterior-medial entorhinal seed region functional activity for scene information was significantly higher than for object information (p < .001), while in the remaining three entorhinal seed regions no significant difference between object and scene conditions existed (see Figure 3).

**Figure 3.**
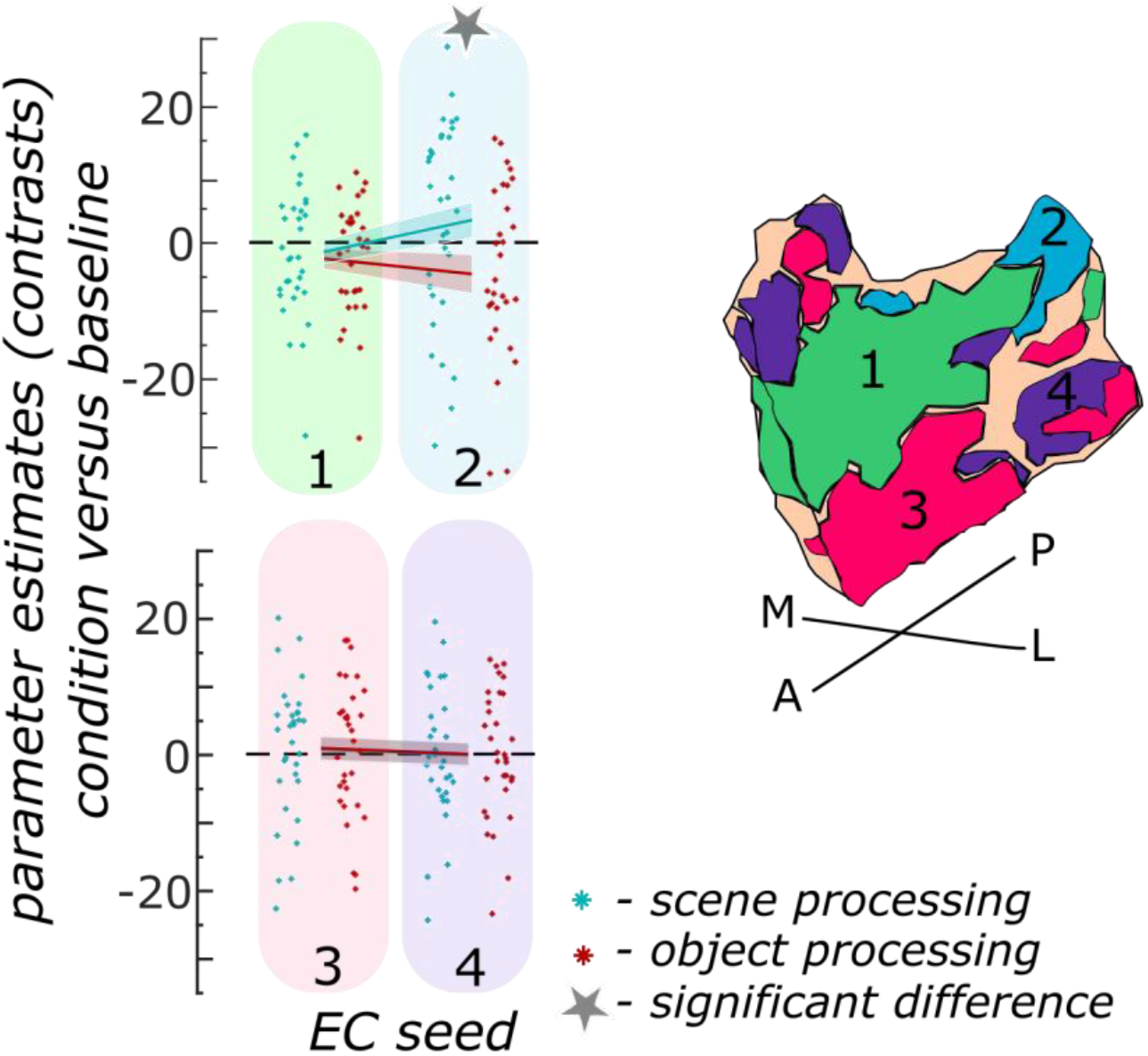
Object and scene information processing in entorhinal seed regions. Displayed are the extracted parameter estimates for the object versus baseline contrast (“item processing”, red) and the scene versus baseline contrast (“contextual processing”, cyan) from each entorhinal seed region per individual (dots) and summarized across individuals (lines). A schematic depiction of the respective entorhinal seed regions is displayed by a 3D drawing of the right entorhinal cortex. A repeated measures ANOVA revealed a significant interaction between condition and seed region. The displayed significant difference is obtained with FDR-corrected post-hoc tests and refers to p < .05. During the object condition, participants were presented with 3D rendered objects on screen, during the scene condition 3D rendered rooms and during the baseline condition they saw scrambled pictures. The shaded area around the lines refer to standard errors of the mean. EC – entorhinal; M – medial; L – lateral; A – anterior; P – posterior.

In the hippocampal subregions, extracting the task-related parameter estimates for object and scene information processing from proximal and distal segments within each participant, showed a significant interaction between transversal segments and information type in the subiculum (F(4,124) = 15.994; p <.001), not however in CA1, as revealed by a repeated measures ANOVA. Post-hoc t-tests revealed that only in the distal subiculum segments significantly more scene than object information was processed (both p < .001). In all other segments along the transversal axis, no significant difference in functional activity related to object and scene processing existed (see Figure 4).

**Figure 4.**
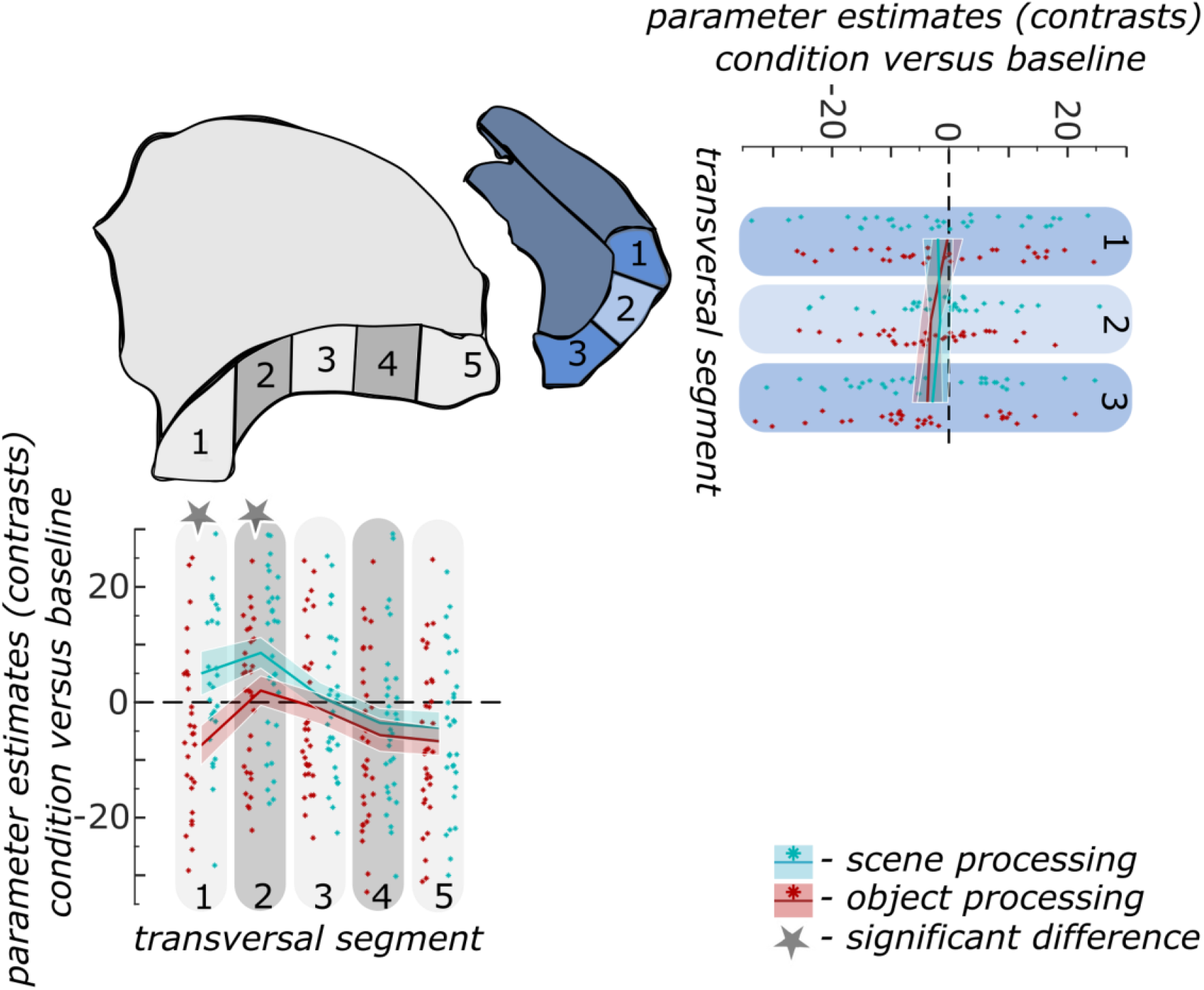
Object and scene information processing along the subiculum and CA1 transversal axis. Displayed are the extracted parameter estimates for the object versus baseline contrast (“item processing”, red) and the scene versus baseline contrast (“contextual processing”, cyan) from the respective transversal segments in the subiculum (grey) and CA1 (blue) per individual (dots) and summarized across individuals (lines). A schematic depiction of the respective transversal segment is displayed by a 3D drawing of the right subiculum and CA1 subregion. Repeated measures ANOVAs revealed a significant interaction between condition and seed region in the subiculum only. The displayed significant difference is obtained with FDR-corrected post-hoc tests and refers to p < .05. During the object condition, participants were presented with 3D rendered objects on screen, during the scene condition rendered 3D rooms and during the baseline condition they saw scrambled pictures. The shaded area around the lines refer to standard errors of the mean. M – medial; L – lateral; A – anterior; P – posterior.

## Discussion

This study aims to advance insight into the organizational principles of information processing along the transversal entorhinal-hippocampal axis and the circuitry’s embedding in large-scale cortical processing. Leveraging ultra-high field 7 Tesla fMRI, we find a resemblance between the intrinsic functional connectivity pattern and subregional biases in object and scene information processing in the entorhinal-hippocampal circuitry. In the EC, we observe a topographical mapping of functionally and clinically relevant regions of the cortical item and contextual information processing pathways, including the retrosplenial, parahippocampal and perirhinal 35 and 36 cortices. This mapping continues to determine a transversal organization of information pathways in the human entorhinal-hippocampal circuitry. Our results unify previous evidence and exhibit novel features in the human brain that can be a window into the circuitry’s critical role in memory function and decline.

### Contextual information is processed within a posterior-medial EC – distal subiculum route

We identified regions in the entorhinal-hippocampal circuitry that are dedicated to process scene information. These regions consisted of two functionally connected portions: the posterior-medial EC and the distal subiculum. The subiculum showed a transversal gradient in intrinsic functional connectivity with a preference to the posterior-medial EC in its distal portions (of note: the posterior-medial EC was defined by preferential functional connectivity to the parahippocampal cortex). Importantly, the distal subiculum and the posterior-medial EC were the only studied entorhinal-hippocampal subregions that exhibited specific scene information processing.

These findings provide clear evidence for a long hypothesized transversal gradient in contextual information processing within the subiculum. Our data also replicate the earlier functional and structural connectivity reports in humans as well as anatomical findings of a route between posterior-medial EC and distal subiculum (Maass et al., 2015; Syversen et al., 2021; Witter et al., 2000). The spatial or contextual information processing bias has, however, mainly been reported for the EC (in rodents: Neunuebel et al., 2013; Knierim et al., 2014; Keene et al., 2016; in humans: Schultz et al., 2012; Reagh et al., 2014; Navarro Schröder et al., 2015; Berron et al., 2018). Long overlooked in animal studies, the importance of the subiculum as a translator of hippocampal information towards the entorhinal and other cortical structures gets more and more acknowledged (O’Mara, 2006; Roy et al., 2017). We here contribute to the very sparse investigations regarding the nature of information processed along the transversal axis of the subiculum (see Ku et al., 2017)Our observation that the distal subiculum is involved in processing scenes more than objects is in line with previous findings in the human brain. The subiculum in general was associated with scene discrimination (Hodgetts et al., 2017). Especially an area that resembles the pre- or distal subiculum is involved in scene construction (Dalton et al., 2018). Recently, a gradient with coarser voxel-wise autocorrelation signals in the medial hippocampus has been reported, a finding that implies larger representations in the distal subiculum (Bouffard et al., 2021). In the latter two studies, however, the authors did not specifically extract data from the transversal axis of hippocampal subfields. Our joint investigation of functional entorhinal-subiculum connectivity and type of information processing along the full transversal subiculum axis, is the first to show a clear gradient of contextual information towards the distal portion.

### Information processing is consistent with convergence within the anterior entorhinal portions – subiculum/CA1 border route

Our data revealed a route that did not display differences in object and scene information processing. Both, the anterior-lateral entorhinal portion (identified based on preferred communication with Area 35) and the anterior-medial entorhinal portion (identified based on preferred communication with the retrosplenial cortex) exhibited preferential functional connectivity with the subiculum/CA1 border. Similar levels of functional activity for object and scene processing along these entorhinal-hippocampal pathways are consistent with information convergence.

While we again confirm earlier findings, several features in our data are fundamentally novel to the human brain. First, we provide initial human evidence for an information pathway between the anterior-lateral EC and the subiculum/CA1 border. Non-primate and primate anatomical data as well as ex-vivo and in vivo structural connectivity data in humans show the possibility of information flow along that route (Syversen et al., 2021; Witter et al., 2017; Witter & Amaral, 1991, 2020). Our results now underpin a functional relevance of that connection beyond the subiculum (for the subiculum see Maass, Berron et al., 2015). Moreover, our findings are derived based on a voxel-wise analysis, unconstrained by a priori selection of regions-of-interest. We thereby confirm the long-held proposal of a transversal information pathway in hippocampal subfields subiculum and CA1.

Convergence of item and contextual information is compatible with recent rodent work that shows joined coding of spatial (location) and non-spatial (object) information along CA1 and within the lateral EC (Deshmukh, 2014; Doan et al., 2019; Vandrey et al., 2021; Wilson et al., 2013; Yeung et al., 2019). However, we cannot confirm reports about higher functional activity for object than scene processing within these areas in the human brain (Reagh and Yassa, 2014; Navarro Schröder et al., 2015; Berron et al., 2018; also indicated in Dalton et al., 2018 and Schultz et al., 2012). Also, we did not observe proximodistal differences in CA1 for non-spatial versus spatial information processing as described in rodents (Beer et al., 2018; Henriksen et al., 2010; Nakamura et al., 2013; Nakazawa et al., 2016). Differences in experimental design and contrasts could have contributed to these discrepancies (i.e. specific item processing versus convergence). Previous studies used a variety of different conditions to tackle item and contextual information processing (e.g. time versus space in Beer et al., 2018 or imagined objects on a 2D or 3D grid Dalton et al., 2018). Some studies performed direct condition contrasts, without comparison to a baseline (e.g. Berron et al., 2018). In contrast to the current data, previous human studies moreover did not derive functional data from specific, functionally defined entorhinal portions in the same dataset, which may have altered the extracted measures.

We observed a previously unreported overlap in functional connectivity profiles of the lateral and medial EC portions in the anterior EC. The sources of these entorhinal portions are the perirhinal Area 35 and the retrosplenial cortex which are part of cortical item and contextual processing streams, respectively. To our knowledge, the retrosplenial-related anterior-medial entorhinal portion has not yet been identified in earlier investigations but anatomical projections from the retrosplenial to deep medial EC layers are confirmed in rodents (Czajkowski et al., 2013; Sugar et al., 2011). Very recently, Syversen et al. (2021) found structural connectivity between the retrosplenial cortex and the medial EC, but not in the anterior part of the medial EC. Their EC segmentation, however, followed different rules which may have contributed to differences in the topographical evaluation of the region. Also, structural and functional connectivity methods may yield different results, in particular as we identified EC subdivisions by another set of contrast regions. The mapping of the anterior-medial EC (identified by retrosplenial connectivity) to the subiculum/CA1 border opposes conventional views that the medial EC communicates with the distal subiculum and proximal CA1 (based on rodent anatomy – see e.g. Nilssen et al., 2019). It is feasible that complex interactions within the EC underlie this observation. First, deep EC layers (that receive retrosplenial input) exchange information with superficial EC layers (that project to the hippocampus (Czajkowski et al., 2013)). Second, cross-projections between the medial and lateral EC exist (Nilssen et al., 2019). Finally, in the primate, an additional longitudinal gradient in EC projections leads to anterior EC connectivity with the subiculum/CA1 border (Witter & Amaral, 2020). The interplay of these complex projections within the EC can be captured in future investigations.

**Figure 4.**
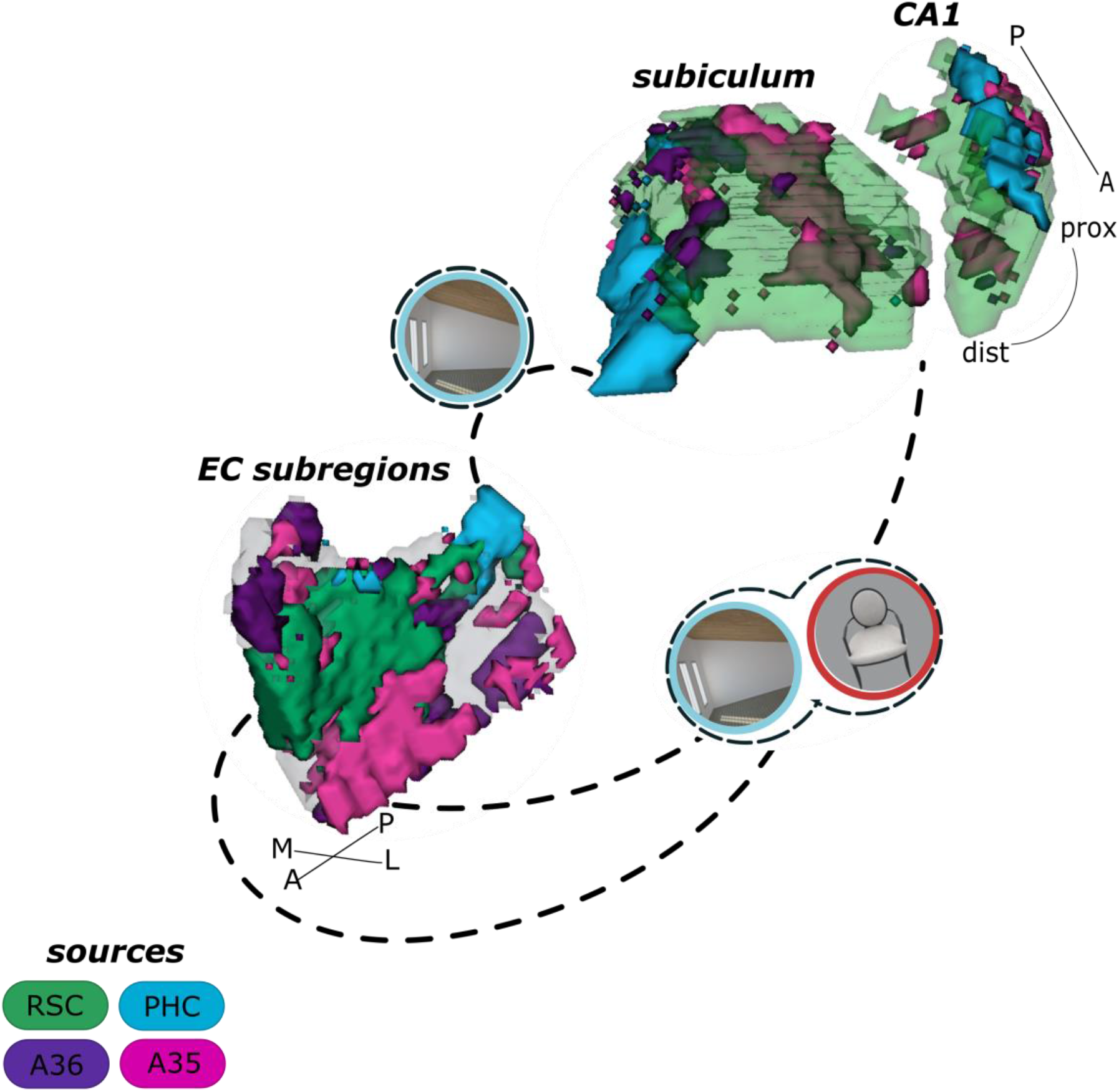
Summary of current results on the information pathways within the entorhinal-hippocampal circuitry. Displayed is a schematic overview of our results on the information pathways within the entorhinal-hippocampal circuitry with four entorhinal seed regions and a focus on the transversal axis of the hippocampal subiculum and CA1 subregions. The four entorhinal seed regions are derived from preferential functional connectivity to retrosplenial (RSC, green), parahippocampal (PHC, blue) and perirhinal Area 36 (A36, purple) and Area 35 (A35, pink) sources. Routes of preferred functional connectivity are depicted with dashed lines and the preferred information processed in the connected areas is depicted with symbolizing icons (scene for contextual, blue and object for item, red; stimuli from the task performed by the participants). M – medial; L – lateral; A – anterior; P – posterior; prox – proximal; dist – distal.

### Relevance of the current findings for the functional organization of the entorhinal-hippocampal circuitry

The current findings advance our insight into the organization of the transversal entorhinal-hippocampal axis on multiple levels and contribute to cognitive and clinical research. Recent efforts to understand how the human entorhinal-hippocampal circuitry accomplishes conjunction and segregation of information largely focused on the longitudinal hippocampal axis (Brunec et al., 2018, 2020; Robin & Moscovitch, 2017). The transversal axis has been approached by studies in humans that did not directly relate connectivity findings to information processing and did not assess subfield-specific organization (Vos de Wael et al., 2018; Plachti et al., 2019; Masouleh et al., 2020; Paquola et al., 2020; Bouffard et al., 2021; for an overview see Genon et al., 2021). Dalton & Maguire (2017), however made a relevant proposal based on visual processing pathways and information processing. In correspondence to our results, they proposed the subiculum/CA1 border as a point of convergence between item and contextual information processing streams. While their conclusion was based on direct parahippocampal, retrosplenial and perirhinal connections to the hippocampus, we found that both, the anterior-lateral EC (that is connected to the cortical item processing stream) and the anterior-medial EC (that is connected to the cortical contextual processing stream) show connectivity with the subiculum/CA1 border. Convergence is potentially also achieved via recurrency within the entorhinal-hippocampal system and cortical regions (cf. Koster et al., 2018 for evidence on recurrency). These considerations are an exciting future research avenue and remain speculative based on the current data due to our insufficient temporal resolution. We hypothesize, however, that within this anterior EC – subiculum/CA1 border route, contextual features of items are bound together, reflecting conjunction outside the hippocampus. In addition, the dedicated context processing that we observe along the posterior-medial EC – distal subiculum route, may functionally underpin ideas about a contextual scaffold that the hippocampus uses to incorporate information into meaningful cohesive memory representations (Behrens et al., 2018; Clewett et al., 2019; Robin, 2018; Robin & Olsen, 2019). Altogether, the information pathways indicate that when a memory is to be formed, some degree of convergence happens already before the hippocampus, nevertheless keeping specific aspects of contextual information separated. Thereby, our results support the necessity of functionally assessing the entorhinal-hippocampal circuitry with high spatial resolution and investigate memory function at a subregional level (Lee et al., 2020). The features we identified can inform future hypotheses on how the hippocampus achieves the formation of cohesive representations that serve memory function.

From a clinical perspective, it is remarkable that the current functional connectivity pattern resembles the topology of cortical tau pathology (Lace et al., 2009). In the literature, it is suggested that tau propagates along functional routes within the brain (Franzmeier et al., 2020; Lace et al., 2009; Vogel et al., 2020). Earliest cortical tau pathology accumulates in the perirhinal Area 35 (also called “transentorhinal” region) and the anterior-lateral EC from where it spreads to the subiculum/CA1 border (Adams et al., 2019; Kaufman et al., 2018). This topology thus mirrors those regions that we find biased towards anterior-lateral entorhinal connectivity (Braak & Braak, 1991; Lace et al., 2009; Roussarie et al., 2020). The related information processing pathway seems affected accordingly as reports on a preclinical relationship between tau-pathology and object processing show (Berron et al., 2018, 2019; Harrison et al., 2019; Maass et al., 2019). Given our pattern of information processing that corresponds to object and scene convergence, related cognitive impairment may, however, be more pronounced for object-in-scene conditions. Moreover, the overlapping functional connectivity pattern in the hippocampus with entorhinal portions based on the retrosplenial cortex and Area 35 open up speculation about advanced Alzheimer’s disease pathology progression, given early hypometabolism and cortical tau progression in the retrosplenial cortex and early amyloid in posterior parietal regions (Grothe et al., 2017; Ziontz et al., 2021). The revealed functional connectivity profile and information pathways may thus specify future hypotheses on the propagation of Alzheimer’s pathology and related functional and cognitive impairment.

## Limitations

This study has a number of limitations. First, the biases in seed connectivity in the left hemisphere were generally weaker and proximal CA1 results were less consistent across hemispheres. We intended to conduct all analyses independently for both hemispheres to provide internal replication of our findings, however, whether partially different effects indeed signal a lateralization of the entorhinal-hippocampal organization in humans or whether the task or another parameter influenced these observations, is subject for further research.

Second, our perspective was entirely functional. To what extent there is a correspondence to structural connectivity (Syversen et al., 2021) remains to be determined, considering different experimental task constraints and contrasts. Note also that as a first step towards an understanding of the system’s functional organization and to increase comparability with earlier studies, we assessed information flow within the entorhinal-hippocampal circuitry with univariate methods that averaged the signal over regions of interest. To capture hidden voxel-wise patterns of activity that scale with the processing of certain representations, future studies could examine information pathways with multivariate methods that evaluate informational content in the activity pattern of voxels instead of in an averaged manner (Kragel et al., 2018; Kriegeskorte et al., 2008). Moreover, recent methodological advances can be employed in the future that study functional connectivity based on the underlying content representations between regions (e.g. Coutanche et al., 2020).

In general, the quantification of the transversal connectivity pattern should be considered with some caution from the anatomist’s perspective. The segmentation of hippocampal subregions on functional data is an approximation because the anatomical ground truth cannot be captured by any segmentation protocol (even histological data leads to divergent opinions). This shortcoming is amplified by group comparisons that do not account for participant-specific anatomy. Note moreover, that for now we excluded the head and the tail of the hippocampus in our investigation. The head is highly complex in its subfield topography (Ding and Van Hoesen, 2015; Berron, Vieweg et al., 2017) and prevents clear hypotheses regarding a transversal pattern. For the tail we lack an sestablished segmentation protocol (DeKraker et al., 2018; Flores et al., 2019). In future, advanced segmentation methods and evaluations in the participant-space will improve this issue and reveal the organization in more detail.

## Conclusion

In sum, leveraging ultra-high field functional imaging, we provide a comprehensive in vivo exploration of the functional organization within the human entorhinal and hippocampal subregions and the circuitry’s embedding within cortical information processing streams. Within the entorhinal and hippocampal subiculum, our data partially support a continuation of cortical item and contextual information processing pathways with convergence in anterior and lateral entorhinal portions, proximal subiculum and CA1, while the posterior-medial entorhinal portion and distal subiculum process scene information specifically. Topographically, this organization of information processing overlaps with our identified pattern of functional connectivity. The data yield spatially organized information pathways along the transversal axis of the human entorhinal-hippocampal circuitry. Our high-resolution approach revealed unknown characteristics of the information flow within the human entorhinal-hippocampal circuitry. These aid our understanding of how cortical information comes together and is further communicated within the entorhinal-hippocampal circuitry, underpinning the formation of cohesive memory representations. We provide essential insights for basic and clinical research that we believe to be crucial for the development of future hypotheses on memory function and decline.

## Methods

The current data is part of a larger study that examines exercise effects on cognition. The data that is subject to the current study have been acquired during the baseline measurement before any intervention took place. In the following, we focus on the study setup and methodological aspects of direct relevance for the current questions and data analyses.

### Participants

In total, 32 healthy participants (15 female) with a mean age of 25.5 years (range 19 to 35 years, standard deviation 4.3 years) were included in the current data analyses. All participants were right-handed, finished education on A-level (German Abitur or comparable) and reported absence of any neurological or psychiatric diseases. General exclusion criteria determined by the 7 Tesla MR scanning procedure were applied (e.g. metallic implants, tinnitus, known metabolic disorders). All participants gave informed consent prior to participation and received a monetary compensation. The study received approval by the ethics committee of Otto-von-Guericke University, Magdeburg (Germany).

### Task

While functional images were acquired, participants engaged in a mnemonic discrimination task (see Berron et al., 2018). The item-context task consisted of 64 objects and 64 scenes. In two runs, participants encoded always two stimuli, two 3D rendered objects in the object condition and two 3D rendered rooms in the scene condition and subsequently identified the following two same or similar stimuli as novel or old. Ten scrambled images were presented in blocks at the beginning and end of each run and served as baseline condition. All stimuli were presented for three seconds. In the recognition phase, participants had to respond during that time. Each stimulus was followed by a noise stimulus to prevent after-image and pop-out effects. The short alternating encoding/recognition sequences were embedded in an event-related design.

### Data acquisition

All MRI data was acquired with a 7 Tesla Siemens MR machine (Erlangen, Germany) using a 32-channel head coil. First structural images were obtained. A whole-brain MPRAGE volume was acquired with isotropic voxel size 0.6mm, TR 2500ms; TE 2.8 ms, 288 slices in an interleaved manner (FOV 384 × 384 × 288). Thereafter, a partial structural T2*-weighted volume (TR 8000ms; TE 76 ms, interleaved, 55 slices, FOV 512 × 512 ×55), orientated orthogonal to the main longitudinal hippocampal axis was obtained with a resolution of 0.4 × 0.4mm in-plane and a slice thickness of 1 mm.

The subsequent acquisition of functional data took place in two runs à 14 min (332 volumes each) employing echo-planar imaging (EPI). The volumes were partial (40 slices, TR 2400 ms, TE 22 ms, FOV 216×216×40, interleaved slice acquisition), oriented along the longitudinal axis of the hippocampus.

All EPIs were distortion corrected with a point-spread function method and motion corrected during online reconstruction (Zaitsev et al., 2004).

## Data analyses

### Preprocessing

Preprocessing and statistical modeling of fMRI data was performed with SPM12 (Wellcome Department of Cognitive Neuroscience, University College, London UK (Penny et al., 2011)). The individual functional images were slice time corrected and smoothed with a full-width half-maximum Gaussian kernel of 1.5 mm. To preserve a high level of anatomical specificity, smoothing was performed with a kernel smaller than two times the voxel size. The artifact detection toolbox ARTrepair (Mozes & Whitfield-Gabrieli, 2011) was subsequently used to identify outliers regarding mean image intensity and motion between scans (threshold in global intensity: 1.3 %; movement threshold: 0.3 mm). Identified outliers are included as spike regressors in subsequent statistical modeling.

Task effects in the functional data were removed by fitting general linear models (with regressors for all task conditions, outliers and movement parameters) to the data. The obtained residual images were saved for the intrinsic functional connectivity analyses later. Note that task-related parameter estimates were extracted for the final information processing analysis, as described later.

### Structural data processing and segmentation

#### Structural template calculation (T1-weighted) and segmentation

To examine and illustrate group-level results later on, a group specific T1-weighted template was calculated using ANTS buildtemplateparallel.sh (Avants et al., 2010). For illustration purposes and to aid group analyses, in addition, the T1 template was manually segmented into hippocampal subregions subiculum and CA1 with ITK-SNAP (Yushkevich et al., 2006) based on the segmentation rules described in Berron, Vieweg et al. (2017). The first slice in each hemisphere that did not contain the uncus anymore, served as start of the hippocampal body in all hippocampal subregions. Moreover, to evaluate results across the transversal axis, the subiculum masks in each hemisphere were sagitally cut in five equally wide segments within each coronal image. As the CA1 region gets more and more tilted towards the hippocampal tail, the three transversal CA1 segments were determined based on manual segmentation. Therefore, the two outer CA1 borders in transversal axis were connected with a line. From the middle point of that line, two straight lines were drawn in a 60° angle to determine equally sized transversal CA1 segments within each coronal slice and hemisphere. Related to the overall size of the subregions, we opted to build five subiculum and three CA1 segments along the transversal axis from proximal to distal ends.

#### Segmentation of individual regions of interest

We manually segmented regions of interest (ROI) in the medial temporal lobe according to the segmentation protocol by Berron, Vieweg et al. (2017). Based on individual t1-weighted and t2-weighted images, the parahippocampal cortex, Area 35, Area 36, the EC and hippocampal subfields subiculum and CA1 are delineated. Moreover, we ran a Freesurfer 6.0 segmentation on the group T1 template to segment the isthmus cingulate cortex as retrosplenial mask (Desikan et al., 2006; Fischl, 2012). Note here that Syversen et al. (2021) used a similar region, however excluded the most superior part. For individual retrosplenial masks, the obtained mask was co-registered from the group T1 template space to the individual t1 space by making use of the alignment matrices obtained during above described T1 group template calculation. For this alignment process we used ANTS WarpImageMultiTransform.sh (Avants et al., 2011). The retrosplenial, parahippocampal and perirhinal Area 36 and Area 35 regions served as source regions for an initial functional connectivity analysis that we conducted to obtain functional subregions within the entorhinal cortex (see upcoming paragraphs and supplementary material II).

#### Co-registration of individual structural data to functional data space

For later functional data extraction, the individual t1-weighted and t2-weighted structural images were co-registered and resliced to the echo-planar images. Therefore, ANTS was used to transfer the t2-weighted structural image to the participant’s t1 space (Avants et al., 2011). For the co-registration between individual t1-weighted and echo-planar images, FSL epi_reg was applied (Jenkinson & Smith, 2001). All subsequently segmented individual masks were co-registered to the participant’s functional (echo-planar) images using the obtained warping matrices. ANTS WarpImageMultiTransform.sh was applied for t2 to t1 co-registration and FSL flirt was used for t1 to echo-planar image co-registration (Avants et al., 2011; Jenkinson & Smith, 2001).

#### ROI preparation for seed regions in functional connectivity analyses

All masks that served as source and seed regions throughout the functional connectivity analyses (retrosplenial, parahippocampal, perirhinal Area 36 and Area 35 and the later defined entorhinal subregions) were thresholded according to mean intensity to prevent signal dropout and thus a distortion of the average functional signal extracted from seed regions for the connectivity analysis. Therefore, we followed Libby et al. (2012) and Maass, Berron et al. (2015), to remove all voxels from each ROI that showed a mean intensity over time of less than two standard deviations from the mean intensity across all voxels. The thresholding was performed before each seed-to-voxel functional connectivity analysis.

### Functional connectivity analyses at the participant level

Two different functional connectivity analyses were performed that build upon the approach by Maass, Berron et al. (2015). The first analysis served to identify functional subregions (“seeds”) within the entorhinal cortex that uniquely connect to functionally and clinically relevant cortical sources. The second, core analysis, then evaluated the intrinsic functional connectivity pattern between these entorhinal seeds and hippocampal subiculum and CA1. In the following we describe the analysis procedure in detail. Note, that all analyses were conducted independently in both hemispheres.

To determine functional entorhinal seed regions we first performed a seed-to-voxel semipartial correlation analysis (Whitfield-Gabrieli & Nieto-Castanon, 2012) between the individually extracted residuals from retrosplenial, parahippocampal and perirhinal Area 36 and Area 35 sources as well as entorhinal voxels. The regions we call cortical sources served as seeds in that analysis. Note that the semipartial correlations calculate the variance in a voxel that is uniquely explained by the source, excluding contributions from other sources. Please refer to supplementary II for more details on this functional semipartial correlation analysis. To obtain entorhinal seeds for the core functional connectivity analysis, first, we calculated one-sample t-tests across participants on the individually obtained and aligned, standardized beta maps for each source, respectively. Second, the four resulting statistical maps (one for each source) have been thresholded at T > 3.1. Each entorhinal voxel now was attributed to be preferentially connected to one of the four source regions, based on the voxel’s maximum T value across the thresholded one-sample t-test maps. Those voxels that did not reach the threshold of T > 3.1 in any of the four statistical maps have not been attributed to be preferentially connected to any of the four cortical sources. Finally, across hemispheres we selected for each source preference an equal number of these highest preference voxels across all t-tests (the number is determined by the hemisphere with the lowest relevant number of voxels). This procedure yielded four entorhinal subregions, one containing the entorhinal voxels that preferentially functionally connect to the retrosplenial (1530 voxels), one containing the entorhinal voxels that preferentially functionally connect to the parahippocampal cortex (145 voxels) and one each that contained the preferentially functionally connected voxels to perirhinal Area 35 (298 voxels) and Area 35 (751 voxels), respectively. All four entorhinal seed masks were determined on group level and co-registered to each participant, then serving as seed regions for the core functional connectivity analysis between entorhinal cortex seeds and hippocampal subregions.

For the core functional connectivity analysis (entorhinal seeds-to-hippocampal subregion voxels), an analogous seed-to-voxel semipartial correlation analysis was performed on the individual residual functional imaging data using the CONN toolbox (Whitfield-Gabrieli & Nieto-Castanon, 2012). Note again that the semipartial correlations calculate the variance in a voxel that is uniquely explained by the seed, excluding contributions from other seeds. Now the four entorhinal subregions served as seeds and functional connectivity was examined with the whole brain (later masked by the hippocampal subregion masks). For each functional connectivity analysis mean time series were extracted from the respective seed region and entered as regressor of interest. White matter and CSF time series, realignment parameters and outliers served as regressor of no interest. The functional data from the residuals was band-pass filtered (0.01-0.1 Hz) and semipartial correlations were obtained between the seed timeseries and all other brain voxel’s timeseries. The obtained beta maps contained Fisher-transformed correlation coefficients and were used for subsequent group analyses.

### Alignment between participants

To be able to perform group statistics on the resulting topography beta maps, the individual data was aligned to group space. Here, the T1 template image served as reference space. Using the inverse of the previously obtained individual warping matrices from EPI to individual t1, first the standardized beta maps were co-registered from epi to individual t1 space. In a further step, the statistical maps were then aligned between individual t1 space to the group T1 template space, by making use of the alignment matrices obtained during above described T1 group template calculation. For this alignment process we used ANTS WarpImageMultiTransform.sh (Avants et al., 2011).

### Functional connectivity analysis at group level

To investigate the functional connectivity profile between the four entorhinal seeds and the subiculum and CA1 subregion across individuals, we evaluated connectivity preferences to either seed within all transversal segments of the subiculum and CA1 target regions. Therefore, mean values for connectivity estimates to either entorhinal cortex seed are extracted from the group aligned but participant-specific beta maps out of each transversal segment, averaged along all coronal slices. Note, that segment-based extraction is necessary due to the varying number of sagittal slices that cover the respective regions along the longitudinal axis of the hippocampal body. Based on these participant-level connectivity results, connectivity preference plots for all four entorhinal seeds have been created to depict tendencies along the transversal axis of the subiculum and subregion CA1.

A hierarchical repeated-measures ANOVA testing procedure was employed to reveal significant differences in the transversal hippocampal connectivity patterns between entorhinal seed regions. Therefore, in a first step, an overall repeated measures ANOVA (4 seed X transversal segments) was performed per target region (subiculum and CA1 in both hemispheres) to reveal whether significant differences in seed connectivity estimates exist across the transversal axis of the respective target region. If the overall seed X transversal segment interaction effect was significant (false-discovery-rate corrected according to Benjamini and Hochberg, 1995), in a second step, one-way repeated measures ANOVAs have been performed for each seed to identify those entorhinal seeds that indeed show a differential connectivity pattern across the transversal axis of the target region (all false-discovery-rate corrected according to Benjamini and Hochberg, 1995). If more than one seed main effect was significant, finally we determined whether these seeds exhibit an opposing connectivity pattern across the transversal axis of the respective hippocampal subregion by evaluating the pair-wise seed X transversal segment interaction effects on the extracted connectivity estimates.

For a more detailed topographical display of the entorhinal-hippocampal connectivity results, we calculated one-sample t-test on the aligned, standardized beta maps that we obtained in the first-level analyses for each seed respectively. Crucially, the resulting group-level one-sample t-test statistical maps were only used to display results but not for any further statistical inference. To depict the topography of the respective voxel-wise seed preferences, the resulting group-level t-maps were thresholded with T> | 0.001| and masked with the respective subregion of interest. To depict general tendencies in the connectivity profile, for each voxel in the region of interest the preferred seed connectivity was determined by attributing it to the seed with the highest T value across the one-sample t-test maps. The resulting maps were depicted in 3D plots, generated with ITK-SNAP (Yushkevich et al., 2006) that provide an overview of each voxel’s preference for the respective seed functional connectivity at a glance.

### Functional analysis of content-related activity at participant level

To investigate whether object and scene information is differentially processed within entorhinal seed regions and along the transversal axis of the subiculum and CA1 body, the results from the initially fitted general linear models (used to remove task effects) were examined. Contrast estimates were calculated between the beta estimates obtained from task conditions in which individuals saw objects versus scenes (rooms) on the screen and conditions in which individuals saw the scrambled stimuli (baseline). The resulting contrast value maps for object > baseline and scene > baseline were then co-registered to the T1 group template space. Subsequently, individual mean contrast estimates have been extracted from the four entorhinal seed regions and from those transversal segments that had previously been used for the evaluation of the intrinsic functional connectivity results (three or five segments in CA1 and subiculum, respectively).

With repeated measures ANOVAs (content condition X entorhinal region or content condition X transversal hippocampal segment) we investigated whether contrast estimates differed for the processing of object versus scene information in the respective regions. Effect of interest thus, was the interaction between the content condition and the subregion or segment, respectively. Post-hoc paired-samples t-test were performed if the respective interaction effect was significant, to reveal in which subregion or segment functional activity between scene and object processing conditions differed significantly from each other (all false-discovery-rate corrected according to Benjamini and Hochberg, 1995).

## Acknowledgements

We thank the Leibniz Institute for Neurobiology in Magdeburg for providing access to the 7 Tesla MRI Scanner. We are grateful for the support of Anne Hochkeppler and Regina Schwarzer with manual segmentations of the medial temporal lobe subregions and for insightful discussions regarding functional connectivity with Yi Chen.

## References

Adams, J. N., Maass, A., Harrison, T. M., Baker, S. L., & Jagust, W. J. (2019). Cortical tau deposition follows patterns of entorhinal functional connectivity in aging. ELife, 8. https://doi.org/10.7554/eLife.49132

Avants, B. B., Tustison, N. J., Song, G., Cook, P. A., Klein, A., & Gee, J. C. (2011). A reproducible evaluation of ANTs similarity metric performance in brain image registration. NeuroImage, 54(3), 2033–2044. https://doi.org/10.1016/j.neuroimage.2010.09.025

Avants, B. B., Yushkevich, P., Pluta, J., Minkoff, D., Korczykowski, M., Detre, J., & Gee, J. C. (2010). The optimal template effect in hippocampus studies of diseased populations. NeuroImage. https://doi.org/10.1016/j.neuroimage.2009.09.062

Beer, Z., Vavra, P., Atucha, E., Rentzing, K., Heinze, H.-J., & Sauvage, M. M. (2018). The memory for time and space differentially engages the proximal and distal parts of the hippocampal subfields CA1 and CA3. PLOS Biology, 16(8), e2006100. https://doi.org/10.1371/journal.pbio.2006100

Behrens, T. E. J., Muller, T. H., Whittington, J. C. R., Mark, S., Baram, A. B., Stachenfeld, K. L., & Kurth-Nelson, Z. (2018). What Is a Cognitive Map? Organizing Knowledge for Flexible Behavior. Neuron, 100(2), 490–509. https://doi.org/10.1016/J.NEURON.2018.10.002

Benjamini, Y., & Hochberg, Y. (1995). Controlling the False Discovery Rate: A Practical and Powerful Approach to Multiple Testing. Journal of the Royal Statistical Society: Series B (Methodological), 57(1), 289–300. https://doi.org/10.1111/j.2517-6161.1995.tb02031.x

Berron, D., Vieweg, P., Hochkeppler, A., Pluta, J. B., Ding, S. L., Maass, A., Luther, A., Xie, L., Das, S. R., Wolk, D. A., Wolbers, T., Yushkevich, P. A., Düzel, E., & Wisse, L. E. M. (2017). A protocol for manual segmentation of medial temporal lobe subregions in 7 Tesla MRI. NeuroImage: Clinical, 15, 466–482. https://doi.org/10.1016/j.nicl.2017.05.022

Berron, David, Cardenas-Blanco, A., Bittner, D., Metzger, C. D., Spottke, A., Heneka, M. T., Fliessbach, K., Schneider, A., Teipel, S. J., Wagner, M., Speck, O., Jessen, F., & Düzel, E. (2019). Higher CSF Tau Levels Are Related to Hippocampal Hyperactivity and Object Mnemonic Discrimination in Older Adults. The Journal of Neuroscience : The Official Journal of the Society for Neuroscience, 39(44), 8788–8797. https://doi.org/10.1523/JNEUROSCI.1279-19.2019

Berron, David, Neumann, K., Maass, A., Schütze, H., Fliessbach, K., Kiven, V., Jessen, F., Sauvage, M., Kumaran, D., & Düzel, E. (2018). Age-related functional changes in domain-specific medial temporal lobe pathways. Neurobiology of Aging, 65, 86–97. https://doi.org/10.1016/J.NEUROBIOLAGING.2017.12.030

Berron, David, van Westen, D., Ossenkoppele, R., Strandberg, O., Hansson, O., Westen, D. van, Ossenkoppele, R., Strandberg, O., Hansson, O., van Westen, D., Ossenkoppele, R., Strandberg, O., & Hansson, O. (2020). Medial temporal lobe connectivity and its associations with cognition in early Alzheimer’s disease. Brain : A Journal of Neurology, 143(4), 1233–1248. https://doi.org/10.1093/brain/awaa068

Berron, David, Vogel, J. W., Insel, P. S., Pereira, J. B., Xie, L., Wisse, L. E. M., Yushkevich, P. A., Palmqvist, S., Mattsson-Carlgren, N., Stomrud, E., Smith, R., Strandberg, O., & Hansson, O. (2021). Early stages of tau pathology and its associations with functional connectivity, atrophy and memory. Brain. https://doi.org/10.1093/brain/awab114

Bouffard, N. R., Golestani, A., Brunec, I. K., Bellana, B., Barense, M. D., & Moscovitch, M. (2021). Single voxel autocorrelation uncovers gradients of temporal dynamics in the hippocampus and entorhinal cortex during rest and navigation. BioRxiv, 2021.07.28.454036. https://doi.org/10.1101/2021.07.28.454036

Braak, H., & Braak, E. (1991). Neuropathological stageing of Alzheimer-related changes. Acta Neuropathologica, 82(4), 239–259. https://doi.org/10.1007/BF00308809

Brunec, I. K., Bellana, B., Ozubko, J. D., Man, V., Robin, J., Liu, Z.-X. X., Grady, C., Rosenbaum, R. S., Winocur, G., Barense, M. D., & Moscovitch, M. (2018). Multiple Scales of Representation along the Hippocampal Anteroposterior Axis in Humans. Current Biology, 28(13), 2129–2135.e6. https://doi.org/10.1016/j.cub.2018.05.016

Brunec, I. K., Robin, J., Olsen, R. K., Moscovitch, M., & Barense, M. D. (2020). Integration and differentiation of hippocampal memory traces. Neuroscience & Biobehavioral Reviews, 118, 196–208. https://doi.org/10.1016/J.NEUBIOREV.2020.07.024

Cembrowski, M. S., Phillips, M. G., DiLisio, S. F., Shields, B. C., Winnubst, J., Chandrashekar, J., Bas, E., & Spruston, N. (2018). Dissociable Structural and Functional Hippocampal Outputs via Distinct Subiculum Cell Classes. Cell, 173(5), 1280–1292.e18. https://doi.org/10.1016/J.CELL.2018.03.031

Clewett, D., DuBrow, S., & Davachi, L. (2019). Transcending time in the brain: How event memories are constructed from experience. Hippocampus, 29(3), 162–183. https://doi.org/10.1002/hipo.23074

Integration of objects and space in perception and memory, 20 Nature Neuroscience 1493 (2017). https://doi.org/10.1038/nn.4657

Coutanche, M. N., Akpan, E., & Buckser, R. R. (2020). Representational Connectivity Analysis: Identifying Networks of Shared Changes in Representational Strength through Jackknife Resampling. BioRxiv, 2020.05.28.103077. https://doi.org/10.1101/2020.05.28.103077

Czajkowski, R., Sugar, J., Zhang, S.-J., Couey, J. J., Ye, J., & Witter, M. P. (2013). Superficially Projecting Principal Neurons in Layer V of Medial Entorhinal Cortex in the Rat Receive Excitatory Retrosplenial Input. Journal of Neuroscience, 33(40), 15779–15792. https://doi.org/10.1523/JNEUROSCI.2646-13.2013

Dalton, M. A., Zeidman, P., McCormick, C., & Maguire, E. A. (2018). Differentiable Processing of Objects, Associations, and Scenes within the Hippocampus. Journal of Neuroscience, 38(38), 8146–8159. https://doi.org/10.1523/JNEUROSCI.0263-18.2018

DeKraker, J., Ferko, K. M., Lau, J. C., Köhler, S., & Khan, A. R. (2018). Unfolding the hippocampus: An intrinsic coordinate system for subfield segmentations and quantitative mapping. NeuroImage, 167, 408–418. https://doi.org/10.1016/J.NEUROIMAGE.2017.11.054

Deshmukh, S. S. (2014). Spatial and Nonspatial Representations in the Lateral Entorhinal Cortex. In Space, Time and Memory in the Hippocampal Formation (pp. 127–152). Springer Vienna. https://doi.org/10.1007/978-3-7091-1292-2_6

Desikan, R. S., Ségonne, F., Fischl, B., Quinn, B. T., Dickerson, B. C., Blacker, D., Buckner, R. L., Dale, A. M., Maguire, R. P., Hyman, B. T., Albert, M. S., & Killiany, R. J. (2006). An automated labeling system for subdividing the human cerebral cortex on MRI scans into gyral based regions of interest. NeuroImage, 31(3), 968–980. https://doi.org/10.1016/J.NEUROIMAGE.2006.01.021

Ding, S.-L., & Van Hoesen, G. W. (2015). Organization and detailed parcellation of human hippocampal head and body regions based on a combined analysis of Cyto-and chemoarchitecture. Journal of Comparative Neurology, 523(15), 2233–2253. https://doi.org/10.1002/cne.23786

Doan, T. P., Lagartos-Donate, M. J., Nilssen, E. S., Ohara, S., & Witter, M. P. (2019). Convergent Projections from Perirhinal and Postrhinal Cortices Suggest a Multisensory Nature of Lateral, but Not Medial, Entorhinal Cortex. Cell Reports, 29(3), 617–627.e7. https://doi.org/10.1016/j.celrep.2019.09.005

Eichenbaum, H., Yonelinas, A. R., & Ranganath, C. (2007). The Medial Temporal Lobe and Recognition Memory. Annual Review of Neuroscience, 30(1), 123–152. https://doi.org/10.1146/annurev.neuro.30.051606.094328

Fischl, B. (2012). FreeSurfer. NeuroImage, 62(2), 774–781. https://doi.org/10.1016/J.NEUROIMAGE.2012.01.021

Flores, R., Berron, D., Ding, S., Ittyerah, R., Pluta, J. B., Xie, L., Adler, D. H., Robinson, J. L., Schuck, T., Trojanowski, J. Q., Grossman, M., Liu, W., Pickup, S., Das, S. R., Wolk, D. A., Yushkevich, P. A., & Wisse, L. E. M. (2019). Characterization of hippocampal subfields using ex vivo MRI and histology data: Lessons for in vivo segmentation. Hippocampus, hipo.23172. https://doi.org/10.1002/hipo.23172

Franzmeier, N., Neitzel, J., Rubinski, A., Smith, R., Strandberg, O., Ossenkoppele, R., Hansson, O., & Ewers, M. (2020). Functional brain architecture is associated with the rate of tau accumulation in Alzheimer’s disease. Nature Communications, 11(1), 347. https://doi.org/10.1038/s41467-019-14159-1

Genon, S., Bernhardt, B. C., La Joie, R., Amunts, K., & Eickhoff, S. B. (2021). The many dimensions of human hippocampal organization and (dys)function. Trends in Neurosciences. https://doi.org/10.1016/J.TINS.2021.10.003

Grothe, M. J., Barthel, H., Sepulcre, J., Dyrba, M., Sabri, O., Teipel, S. J., & Initiative, F. the A. D. N. (2017). In vivo staging of regional amyloid deposition. Neurology, 89(20), 2031–2038. https://doi.org/10.1212/WNL.0000000000004643

Harrison, T. M., Maass, A., Adams, J. N., Du, R., Baker, S. L., & Jagust, W. J. (2019). Tau deposition is associated with functional isolation of the hippocampus in aging. Nature Communications 2019 10:1, 10(1), 1–12. https://doi.org/10.1038/s41467-019-12921-z

Haxby, J. V, Grady, C. L., Horwitz, B., Ungerleider, L. G., Mishkin, M., Carson, R. E., Herscovitch, P., Schapiro, M. B., & Rapoport, S. I. (1991). Dissociation of object and spatial visual processing pathways in human extrastriate cortex. Proceedings of the National Academy of Sciences of the United States of America, 88(5), 1621–1625. https://doi.org/10.1073/pnas.88.5.1621

Henriksen, E. J., Colgin, L. L., Barnes, C. A., Witter, M. P., Moser, M.-B., & Moser, E. I. (2010). Spatial representation along the proximodistal axis of CA1. Neuron, 68(1), 127–137. https://doi.org/10.1016/j.neuron.2010.08.042

Hodgetts, C. J., Voets, N. L., Thomas, A. G., Clare, S., Lawrence, A. D., & Graham, K. S. (2017). Ultra-High-Field fMRI Reveals a Role for the Subiculum in Scene Perceptual Discrimination. The Journal of Neuroscience : The Official Journal of the Society for Neuroscience, 37(12), 3150–3159. https://doi.org/10.1523/JNEUROSCI.3225-16.2017

Functional diversity along the transverse axis of hippocampal area CA1, 588 FEBS Letters 2470 (2014). https://www.sciencedirect.com/science/article/pii/S001457931400444X

Igarashi, K. M., & Moser, E. I. (2015). The entorhinal map of space. Brain Research. https://doi.org/10.1016/j.brainres.2015.10.041

Jacobs, H. I. L., Hedden, T., Schultz, A. P., Sepulcre, J., Perea, R. D., Amariglio, R. E., Papp, K. V., Rentz, D. M., Sperling, R. A., & Johnson, K. A. (2018). Structural tract alterations predict downstream tau accumulation in amyloid-positive older individuals. Nature Neuroscience 2018 21:3, 21(3), 424–431. https://doi.org/10.1038/s41593-018-0070-z

Jenkinson, M., & Smith, S. (2001). A global optimisation method for robust affine registration of brain images. Medical Image Analysis. https://doi.org/10.1016/S1361-8415(01)00036-6

Kaufman, S. K., Del Tredici, K., Thomas, T. L., Braak, H., & Diamond, M. I. (2018). Tau seeding activity begins in the transentorhinal/entorhinal regions and anticipates phospho-tau pathology in Alzheimer’s disease and PART. Acta Neuropathologica, 136(1). https://doi.org/10.1007/s00401-018-1855-6

Keene, C. S., Bladon, J., McKenzie, S., Liu, C. D., O’Keefe, J., & Eichenbaum, H. (2016). Complementary Functional Organization of Neuronal Activity Patterns in the Perirhinal, Lateral Entorhinal, and Medial Entorhinal Cortices. The Journal of Neuroscience : The Official Journal of the Society for Neuroscience, 36(13), 3660–3675. https://doi.org/10.1523/JNEUROSCI.4368-15.2016

Kharabian Masouleh, S., Plachti, A., Hoffstaedter, F., Eickhoff, S., & Genon, S. (2020). Characterizing the gradients of structural covariance in the human hippocampus. NeuroImage, 218, 116972. https://doi.org/10.1016/J.NEUROIMAGE.2020.116972

Knierim, J. J., Neunuebel, J. P., & Deshmukh, S. S. (2014). Functional correlates of the lateral and medial entorhinal cortex: objects, path integration and local-global reference frames. Philosophical Transactions of the Royal Society of London. Series B, Biological Sciences, 369(1635), 20130369. https://doi.org/10.1098/rstb.2013.0369

Koster, R., Chadwick, M. J., Chen, Y., Berron, D., Banino, A., Düzel, E., Hassabis, D., & Kumaran, D. (2018). Big-Loop Recurrence within the Hippocampal System Supports Integration of Information across Episodes. Neuron, 99(6), 1342–1354.e6. https://doi.org/10.1016/J.NEURON.2018.08.009

Kragel, P. A., Koban, L., Barrett, L. F., & Wager, T. D. (2018). Representation, Pattern Information, and Brain Signatures: From Neurons to Neuroimaging. Neuron, 99(2), 257–273. https://doi.org/10.1016/J.NEURON.2018.06.009

Kriegeskorte, N., Mur, M., & Bandettini, P. A. (2008). Representational similarity analysis – connecting the branches of systems neuroscience. Frontiers in Systems Neuroscience, 2, 4. https://doi.org/10.3389/neuro.06.004.2008

Ku, S., Nakamura, N. H., Maingret, N., Mahnke, L., Yoshida, M., & Sauvage, M. M. (2017). Regional Specific Evidence for Memory-Load Dependent Activity in the Dorsal Subiculum and the Lateral Entorhinal Cortex. Frontiers in Systems Neuroscience, 11, 51. https://doi.org/10.3389/fnsys.2017.00051

Lace, G., Savva, G. M., Forster, G., de Silva, R., Brayne, C., Matthews, F. E., Barclay, J. J., Dakin, L., Ince, P. G., Wharton, S. B., & MRC-CFAS, on behalf of. (2009). Hippocampal tau pathology is related to neuroanatomical connections: an ageing population-based study. Brain, 132(5), 1324–1334. https://doi.org/10.1093/brain/awp059

Lee, H., GoodSmith, D., & Knierim, J. J. (2020). Parallel processing streams in the hippocampus. Current Opinion in Neurobiology, 64, 127–134. https://doi.org/10.1016/j.conb.2020.03.004

Libby, L. A., Ekstrom, A. D., Ragland, J. D., & Ranganath, C. (2012). Differential connectivity of perirhinal and parahippocampal cortices within human hippocampal subregions revealed by high-resolution functional imaging. The Journal of Neuroscience : The Official Journal of the Society for Neuroscience, 32(19), 6550–6560. https://doi.org/10.1523/JNEUROSCI.3711-11.2012

Maass, A., Berron, D., Harrison, T. M., Adams, J. N., La Joie, R., Baker, S., Mellinger, T., Bell, R. K., Swinnerton, K., Inglis, B., Rabinovici, G. D., Düzel, E., & Jagust, W. J. (2019). Alzheimer’s pathology targets distinct memory networks in the ageing brain. Brain, 142(8), 2492–2509. https://doi.org/10.1093/brain/awz154

Maass, A., Berron, D., Libby, L. A., Ranganath, C., & Düzel, E. (2015). Functional subregions of the human entorhinal cortex. ELife, 4, e06426. https://doi.org/10.7554/eLife.06426

Nakamura, N. H., Flasbeck, V., Maingret, N., Kitsukawa, T., & Sauvage, M. M. (2013). Proximodistal segregation of nonspatial information in CA3: preferential recruitment of a proximal CA3-distal CA1 network in nonspatial recognition memory. The Journal of Neuroscience : The Official Journal of the Society for Neuroscience, 33(28), 11506–11514. https://doi.org/10.1523/JNEUROSCI.4480-12.2013

Nakazawa, Y., Pevzner, A., Tanaka, K. Z., & Wiltgen, B. J. (2016). Memory retrieval along the proximodistal axis of CA1. Hippocampus, 26(9), 1140–1148. https://doi.org/10.1002/hipo.22596

Navarro Schröder, T., Haak, K. V, Zaragoza Jimenez, N. I., Beckmann, C. F., & Doeller, C. F. (2015). Functional topography of the human entorhinal cortex. ELife, 4. https://doi.org/10.7554/eLife.06738

Neunuebel, J. P., Yoganarasimha, D., Rao, G., & Knierim, J. J. (2013). Conflicts between local and global spatial frameworks dissociate neural representations of the lateral and medial entorhinal cortex. The Journal of Neuroscience : The Official Journal of the Society for Neuroscience, 33(22), 9246–9258. https://doi.org/10.1523/JNEUROSCI.0946-13.2013

Nilssen, E. S., Doan, T. P., Nigro, M. J., Ohara, S., & Witter, M. P. (2019). Neurons and networks in the entorhinal cortex: A reappraisal of the lateral and medial entorhinal subdivisions mediating parallel cortical pathways. Hippocampus, 29(12), 1238–1254. https://doi.org/10.1002/hipo.23145

O’Mara, S. (2006). Controlling hippocampal output: the central role of subiculum in hippocampal information processing. Behavioural Brain Research, 174(2), 304–312. https://doi.org/10.1016/J.BBR.2006.08.018

Paquola, C., Benkarim, O., DeKraker, J., Larivière, S., Frässle, S., Royer, J., Tavakol, S., Valk, S., Bernasconi, A., Bernasconi, N., Khan, A., Evans, A. C., Razi, A., Smallwood, J., & Bernhardt, B. C. (2020). Convergence of cortical types and functional motifs in the human mesiotemporal lobe. ELife, 9. https://doi.org/10.7554/eLife.60673

Penny, W. D., Friston, K. J., Ashburner, J. T., Kiebel, S. J., & Nichols, T. E. (2011). Statistical parametric mapping: the analysis of functional brain images. Elsevier.

Plachti, A., Eickhoff, S. B., Hoffstaedter, F., Patil, K. R., Laird, A. R., Fox, P. T., Amunts, K., & Genon, S. (2019). Multimodal Parcellations and Extensive Behavioral Profiling Tackling the Hippocampus Gradient. Cerebral Cortex, 29(11), 4595–4612. https://doi.org/10.1093/cercor/bhy336

Ranganath, C., & Ritchey, M. (2012). Two cortical systems for memory-guided behaviour. Nature Reviews Neuroscience, 13(10), 713–726. https://doi.org/10.1038/nrn3338

Reagh, Z. M., Watabe, J., Ly, M., Murray, E., & Yassa, M. A. (2014). Dissociated signals in human dentate gyrus and CA3 predict different facets of recognition memory. The Journal of Neuroscience : The Official Journal of the Society for Neuroscience, 34(40), 13301–13313. https://doi.org/10.1523/JNEUROSCI.2779-14.2014

Reagh, Z. M., & Yassa, M. a. (2014). Object and spatial mnemonic interference differentially engage lateral and medial entorhinal cortex in humans. Proceedings of the National Academy of Sciences of the United States of America, 111(40). https://doi.org/10.1073/pnas.1411250111

Ritchey, M., Libby, L. A., & Ranganath, C. (2015). Cortico-hippocampal systems involved in memory and cognition: The PMAT framework. In Progress in Brain Research (1st ed., Vol. 219). Elsevier B.V. https://doi.org/10.1016/bs.pbr.2015.04.001

Robin, J. (2018). Spatial scaffold effects in event memory and imagination. Wiley Interdisciplinary Reviews: Cognitive Science, 9(4), e1462. https://doi.org/10.1002/WCS.1462

Robin, J., & Moscovitch, M. (2017). Details, gist and schema: hippocampal–neocortical interactions underlying recent and remote episodic and spatial memory. Current Opinion in Behavioral Sciences, 17, 114–123. https://doi.org/10.1016/J.COBEHA.2017.07.016

Robin, J., & Olsen, R. K. (2019). Scenes facilitate associative memory and integration. Learning & Memory (Cold Spring Harbor, N.Y.), 26(7), 252–261. https://doi.org/10.1101/lm.049486.119

Roussarie, J.-P., Yao, V., Rodriguez-Rodriguez, P., Oughtred, R., Rust, J., Plautz, Z., Kasturia, S., Albornoz, C., Wang, W., Schmidt, E. F., Dannenfelser, R., Tadych, A., Brichta, L., Barnea-Cramer, A., Heintz, N., Hof, P. R., Heiman, M., Dolinski, K., Flajolet, M., … Greengard, P. (2020). Selective Neuronal Vulnerability in Alzheimer’s Disease: A Network-Based Analysis. Neuron, 0(0). https://doi.org/10.1016/j.neuron.2020.06.010

Roy, D. S., Kitamura, T., Okuyama, T., Ogawa, S. K., Sun, C., Obata, Y., Yoshiki, A., & Tonegawa, S. (2017). Distinct Neural Circuits for the Formation and Retrieval of Episodic Memories. Cell, 170(5), 1000–1012.e19. https://doi.org/10.1016/j.cell.2017.07.013

Sugar, J., Witter, M. P., van Strien, N. M., & Cappaert, N. L. M. (2011). The retrosplenial cortex: Intrinsic connectivity and connections with the (para)hippocampal region in the rat. An interactive connectome. Frontiers in Neuroinformatics, 5, 7. https://doi.org/10.3389/FNINF.2011.00007/ABSTRACT

Syversen, I. F., Witter, M. P., Kobro-Flatmoen, A., Goa, P. E., Schröder, T. N., & Doeller, C. F. (2021). Structural connectivity-based segmentation of the human entorhinal cortex. BioRxiv, 2021.07.16.452500. https://doi.org/10.1101/2021.07.16.452500

Ungerleider, L. G., & Haxby, J. V. (1994). ‘What’ and ‘where’ in the human brain. Current Opinion in Neurobiology, 4(2), 157–165. https://doi.org/10.1016/0959-4388(94)90066-3

Vandrey, B., Duncan, S., & Ainge, J. A. (2021). Object and object-memory representations across the proximodistal axis of CA1. Hippocampus, hipo.23331. https://doi.org/10.1002/hipo.23331

Vogel, J. W., Iturria-Medina, Y., Strandberg, O. T., Smith, R., Levitis, E., Evans, A. C., & Hansson, O. (2020). Spread of pathological tau proteins through communicating neurons in human Alzheimer’s disease. Nature Communications, 11(1), 2612. https://doi.org/10.1038/s41467-020-15701-2

Vos de Wael, R., Larivière, S., Caldairou, B., Hong, S.-J., Margulies, D. S., Jefferies, E., Bernasconi, A., Smallwood, J., Bernasconi, N., & Bernhardt, B. C. (2018). Anatomical and microstructural determinants of hippocampal subfield functional connectome embedding. Proceedings of the National Academy of Sciences of the United States of America, 115(40), 10154–10159. https://doi.org/10.1073/pnas.1803667115

Whitfield-Gabrieli, S., & Nieto-Castanon, A. (2012). Conn: a functional connectivity toolbox for correlated and anticorrelated brain networks. Brain Connectivity, 2(3), 125–141. https://doi.org/10.1089/brain.2012.0073

Wilson, D. I. G., Langston, R. F., Schlesiger, M. I., Wagner, M., Watanabe, S., & Ainge, J. A. (2013). Lateral entorhinal cortex is critical for novel object-context recognition. Hippocampus, 23(5), 352–366. https://doi.org/10.1002/hipo.22095

Witter, M. P., & Amaral, D. G. (1991). Entorhinal cortex of the monkey: V. Projections to the dentate gyrus, hippocampus, and subicular complex. The Journal of Comparative Neurology, 307(3), 437–459. https://doi.org/10.1002/cne.903070308

Witter, M. P., & Amaral, D. G. (2020). The entorhinal cortex of the monkey: VI. Organization of projections from the hippocampus, subiculum, presubiculum, and parasubiculum. Journal of Comparative Neurology, cne.24983. https://doi.org/10.1002/cne.24983

Witter, M. P., Doan, T. P., Jacobsen, B., Nilssen, E. S., & Ohara, S. (2017). Architecture of the Entorhinal Cortex A Review of Entorhinal Anatomy in Rodents with Some Comparative Notes. Frontiers in Systems Neuroscience, 11, 46. https://doi.org/10.3389/fnsys.2017.00046

Witter, M. P., Naber, P. A., van Haeften, T., Machielsen, W. C. M., Rombouts, S. A. R. B., Barkhof, F., Scheltens, P., & Lopes da Silva, F. H. (2000). Cortico-hippocampal communication by way of parallel parahippocampal-subicular pathways. Hippocampus, 10(4), 398–410. https://doi.org/10.1002/1098-1063(2000)10:4<398::AID-HIPO6>3.0.CO;2-K

Yeung, L.-K., Olsen, R. K., Hong, B., Mihajlovic, V., D’Angelo, M. C., Kacollja, A., Ryan, J. D., & Barense, M. D. (2019). Object-in-place Memory Predicted by Anterolateral Entorhinal Cortex and Parahippocampal Cortex Volume in Older Adults. Journal of Cognitive Neuroscience, 31(5), 711–729. https://doi.org/10.1162/jocn_a_01385

Yushkevich, P. A., Piven, J., Hazlett, H. C., Smith, R. G., Ho, S., Gee, J. C., & Gerig, G. (2006). User-guided 3D active contour segmentation of anatomical structures: Significantly improved efficiency and reliability. NeuroImage, 31(3), 1116–1128. https://doi.org/10.1016/J.NEUROIMAGE.2006.01.015

Zaitsev, M., Hennig, J., & Speck, O. (2004). Point spread function mapping with parallel imaging techniques and high acceleration factors: Fast, robust, and flexible method for echo-planar imaging distortion correction. Magnetic Resonance in Medicine, 52(5), 1156–1166. https://doi.org/10.1002/mrm.20261

Zhang, S.-J., Ye, J., Couey, J. J., Witter, M., Moser, E. I., & Moser, M.-B. (2014). Functional connectivity of the entorhinal–hippocampal space circuit. Philosophical Transactions of the Royal Society B: Biological Sciences, 369(1635). https://doi.org/10.1098/RSTB.2012.0516

Ziontz, J., Adams, J., Harrison, T., Baker, S., & Jagust, W. (2021). Hippocampal connectivity with retrosplenial cortex drives neocortical tau accumulation and memory function. https://doi.org/10.21203/RS.3.RS-235539/V1

