## Supplementary for "Functional connectivity and information pathways in the human entorhinal-hippocampal circuitry"

### I. LEFT HEMISPHERE RESULTS

**Four cortical sources split the left entorhinal cortex (EC) in anterior-lateral, posterior-medial, anterior-medial and posterior-lateral seeds**

In the left hemisphere we obtained a largely posterior-medial entorhinal seed based on functional connectivity preferences to the parahippocampal cortex. Based on preferences to the retrosplenial cortex, we obtained a rather anterior-medial entorhinal seed. In contrast to these medially located entorhinal seeds, we obtained lateral seeds based on connectivity preferences to the perirhinal Area 36 and Area 35. Anterior-lateral entorhinal voxels preferentially connected to Area 35 while posterior-lateral entorhinal voxels preferentially connected to Area 36. Note that both entorhinal seeds extended longitudinally such that the seed based on connectivity with Area 35 progresses more along deep entorhinal portions and the one based on connectivity with Area 36 along superficial entorhinal portions (see Figure S1).

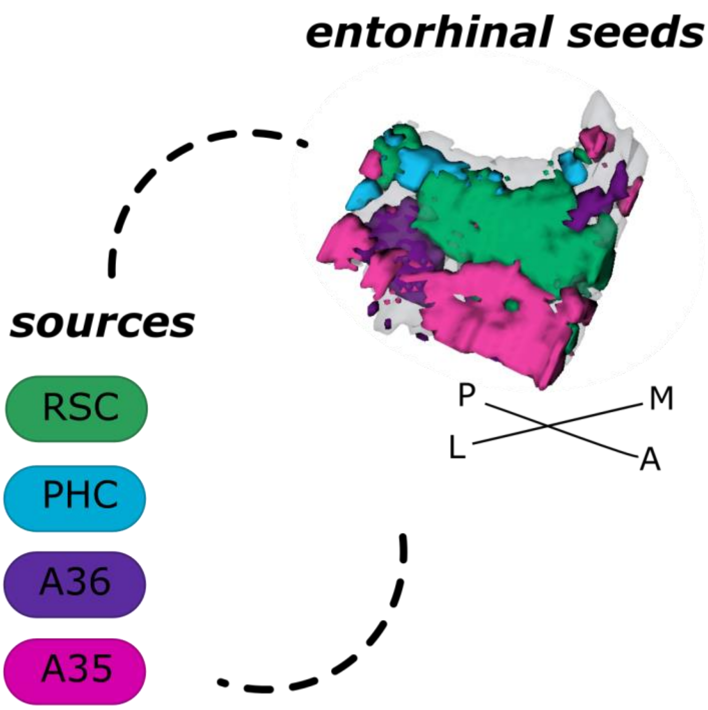


***Figure S1. Left entorhinal seed regions based on connectivity preferences to cortical regions.*** Displayed is the left entorhinal cortex as a 3D image with colored seed regions. The seed regions have been identified based on a source-to-voxel functional connectivity analysis and resulting connectivity preference to either the left retrosplenial (RSC, green) cortex, parahippocampal cortex (PHC, blue), Area 36 (A36, purple) or Area 35 (A35, pink) sources. Seed regions have been determined based on the maximum voxels across four one-sample t-tests at group level, one per source. M – medial; L – lateral; A – anterior; P – posterior.

**The left distal subiculum functionally connects to the posterior-medial EC and the subiculum/CA1 border to anterior-medial and anterior-lateral EC**

When extracting estimates of connectivity preferences across individuals from proximal and distal hippocampal subfield segments for either entorhinal seed, repeated measures ANOVAs revealed significant seed X segments interaction effects along the transversal axis of the left subiculum and CA1 (subiculum: $F(12,372) = 4.609$; $p < .001$; CA1: $F(6,186) = 2.458$; $p = .047$; FDR-corrected; see Figure 2).

In the left subiculum, additional repeated measures ANOVAs showed that the anterior-lateral ($F(4,124) = 4.489$; $p = .025$), and posterior-medial ($F(4,124) = 8.701$; $p < .001$) entorhinal seeds displayed a significant main effect across the transversal subiculum axis. Here, the transversal preference to the anterior-medial entorhinal seed does not survive FDR correction ($F(4,124) = 4.489$; Huyn-Field uncorrected $p = .05$), shows however the same tendency as in the right hemisphere. The differential functional connectivity preferences for the anterior-lateral and posterior-medial entorhinal seed interacted significantly across the transversal axis, as shown in a subsequent repeated measures ANOVA ($F(4,124) = 10.795$; $p < .001$).

In the left CA1, additional repeated measures ANOVAs showed that the connectivity preference towards the anterior-medial entorhinal seed displayed a significant main effect across the transversal CA1 axis ($F(2,62) = 6.753$; $p = .024$; FDR-corrected). In the distal CA1, the preferential functional connectivity with the posterior-medial entorhinal seed was higher than in the proximal portion of CA1. In the left CA1, a similar but weaker transversal pattern was observed for connectivity preferences with the posterior-lateral (marginal significant; $F(2,62) = 3.841$; $p = .051$) and posterior-medial entorhinal seed regions ($F(2,62) = 3.468$; $p = .051$).

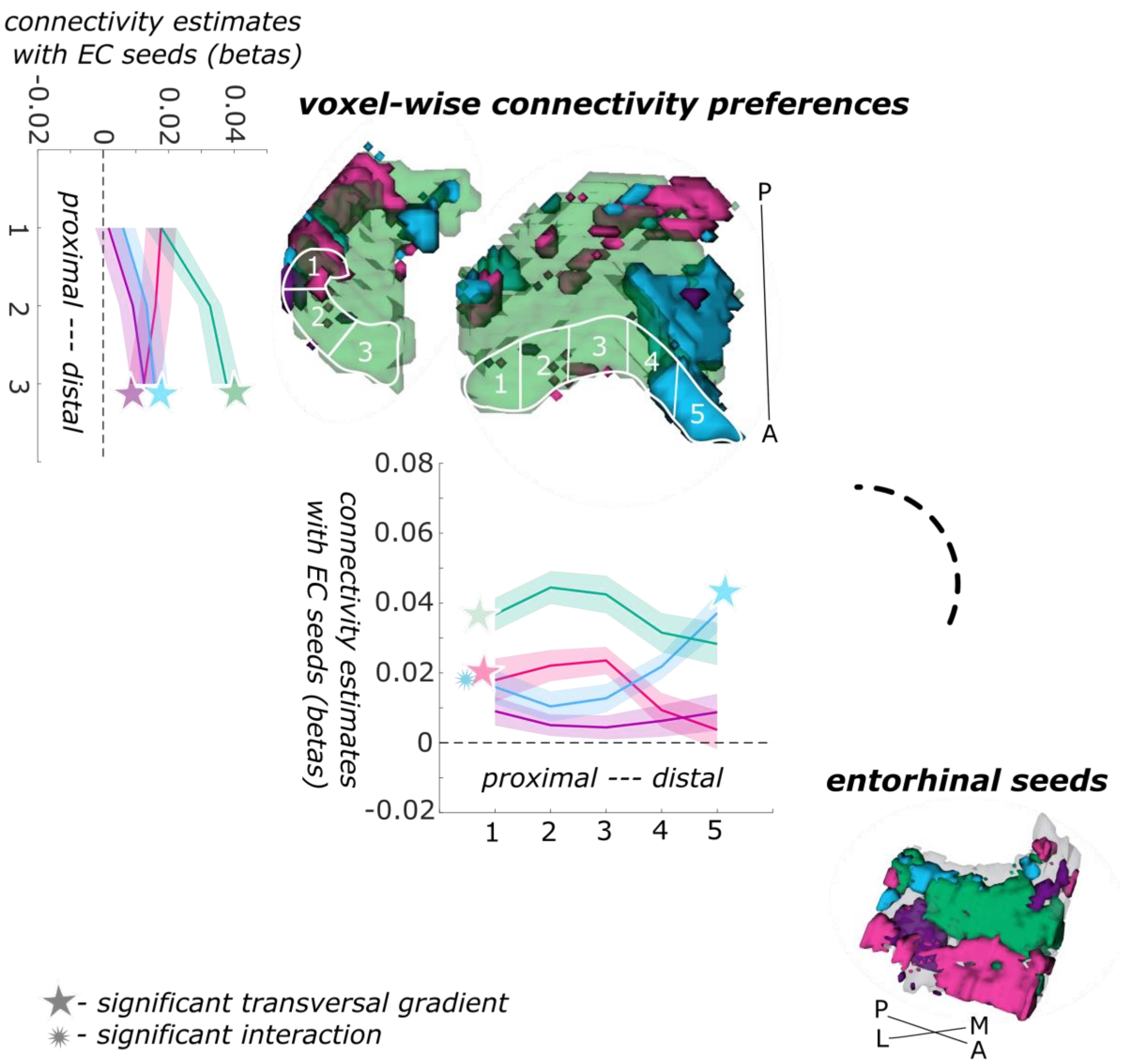


***Figure S2. Functional connectivity preferences to entorhinal seeds along the subiculum and CA1 transversal axis, left hemisphere.*** Displayed are the results of a seed-to-voxel functional connectivity analysis between the displayed left entorhinal seeds and the right subiculum and CA1 subregion. The 3D figure shows voxel-wise connectivity preferences to the entorhinal seeds (color coded to refer to the respective entorhinal seed) on group level. To display mean connectivity preferences across participants along the transversal axis, beta estimates were extracted and averaged from equally sized segments from proximal to distal ends (five segments in subiculum, three segments in CA1; schematized in white on the 3D figures) on each coronal slice and averaged along the longitudinal axis. Repeated measures ANOVAs revealed significant differences in connectivity estimates along the transversal axis in subiculum and CA1 with interaction effects in the subiculum. Displayed significances obtained by FDR-corrected post-hoc tests and refer to $p < .05$. Shaded areas in the graphs refer to standard errors of the mean. EC – entorhinal; M – medial; L – lateral; A – anterior; P – posterior.

**The distal subiculum and posterior-medial EC specifically process scene information while other subregions similarly process object and scene information**

For the characteristics of information processing, we first focus on the left entorhinal seed regions. When extracting task-related parameter estimates for object and scene information processing, a repeated measures ANOVA showed a significant interaction between region and information type (object versus scene; $F(3,93) = 9.772$; $p < .001$). Post-hoc t-tests revealed that only in the posterior-medial entorhinal seed region functional activity for scene information was significantly higher than for object information ($p = .003$; FDR-corrected), while in the remaining three left entorhinal seed regions no significant difference between object and scene conditions existed (see Figure S3).

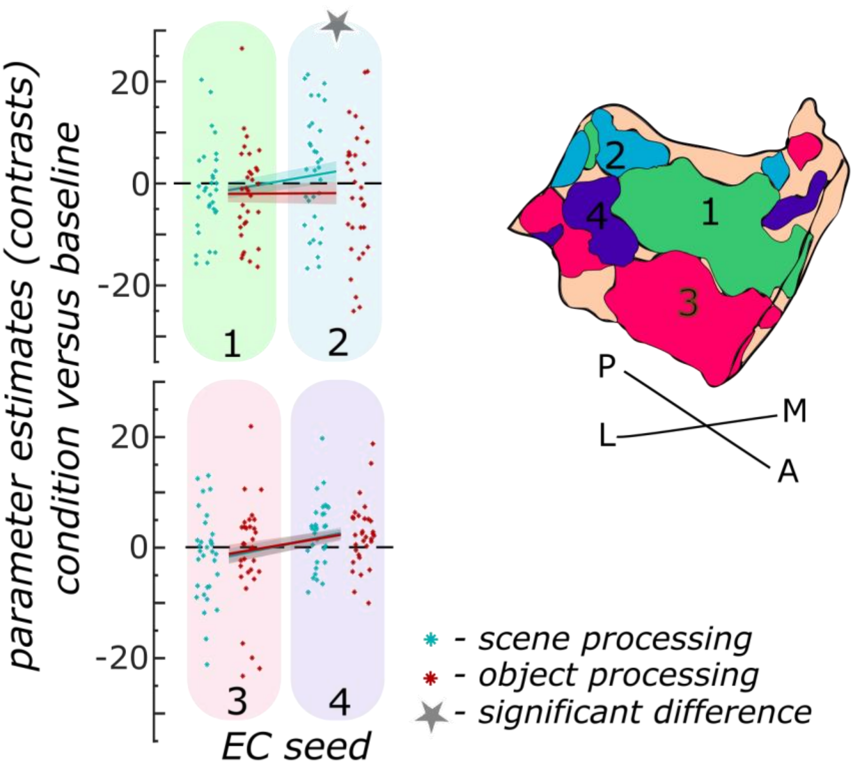


***Figure S3. Object and scene information processing in left entorhinal seed regions.*** Displayed are the extracted parameter estimates for the object versus baseline contrast (red) and the scene versus baseline contrast (cyan) from each left entorhinal seed region per individual (dots) and summarized across individuals (lines). A schematic depiction of the respective entorhinal seed regions is displayed by a 3D drawing of the right EC. A repeated measures ANOVA revealed a significant interaction between condition and seed region. The displayed significant difference is obtained with FDR-corrected post-hoc tests and refers to $p < .05$. During the object condition, participants were presented with 3D rendered objects on screen, during the scene condition with 3D rendered rooms and during the baseline condition they saw scrambled pictures. The shaded area around the lines refer to standard errors of the mean. EC – entorhinal; M – medial; L – lateral; A – anterior; P – posterior.

In the left hippocampal subregions, extracting the task-related parameter estimates for object and scene processing from proximal and distal segments within each participant showed a significant interaction between transversal segments and information type in the subiculum ($F(4,124) = 7.697$; $p < .001$), not however in CA1 as revealed by a repeated measures ANOVA. Post-hoc t-tests showed that only in the distal subiculum segments significantly more scene than object information was processed (both $p < .001$; middle segment uncorrected $p = .02$). In all other segments along the transversal axis, no significant difference in functional activity related to object and scene processing existed (see Figure S4).

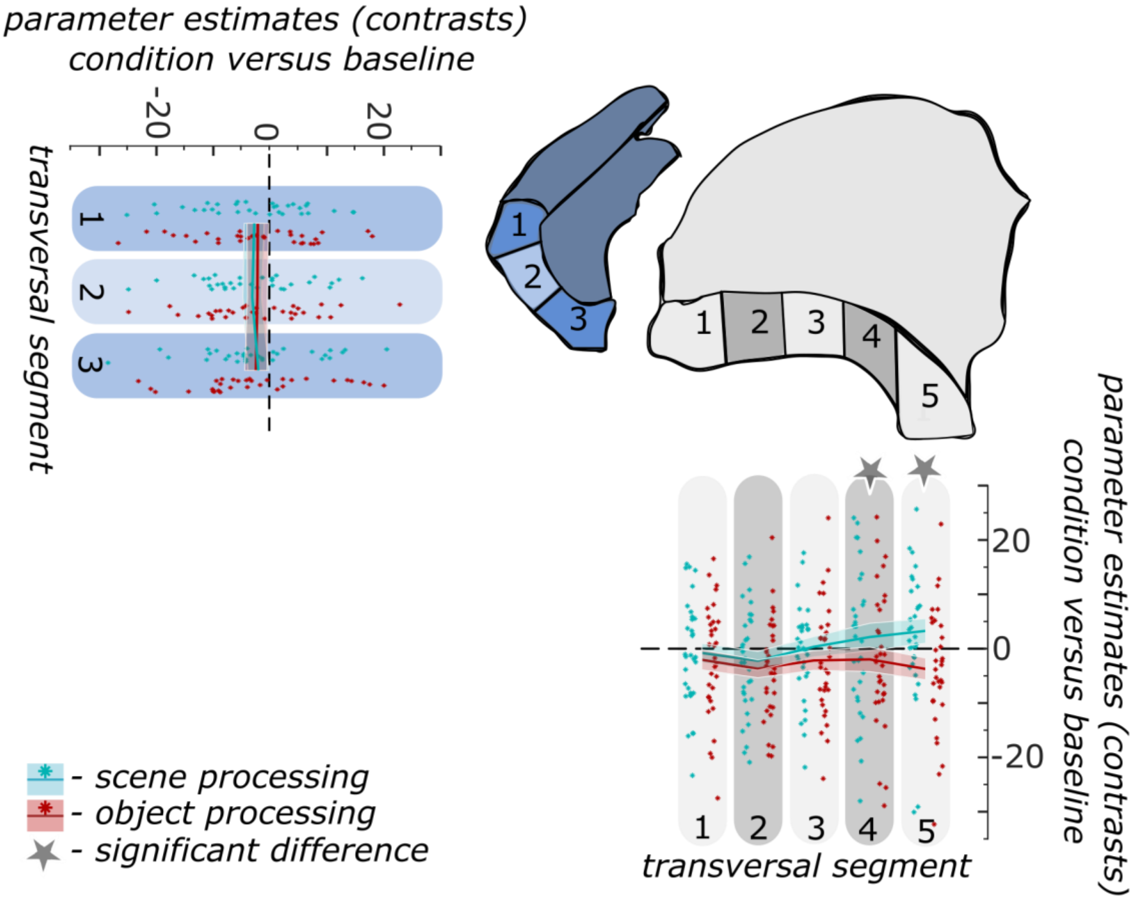


***Figure S4. Object and scene information processing along the left subiculum and CA1 transversal axis.*** Displayed are the extracted parameter estimates for the object versus baseline contrast (red) and the scene versus baseline contrast (cyan) from the respective transversal segments in the subiculum (grey) and CA1 (blue) per individual (dots) and summarized across individuals (lines). A schematic depiction of the respective transversal segment is displayed by a 3D drawing of the right subiculum and CA1 subregions. Repeated measures ANOVAs revealed a significant interaction between condition and seed region in the subiculum only. The displayed significant difference is obtained with FDR-corrected post-hoc tests and refers to $p < .05$. During the object condition, participants were presented with 3D rendered objects on screen, during the scene condition with 3D rendered rooms sand during the baseline condition they saw scrambled pictures. The shaded area around the lines refer to standard errors of the mean. M – medial; L – lateral; A – anterior; P – posterior.

### II. FUNCTIONAL CONNECTIVITY ANALYSIS TO DETERMINE ENTORHINAL SEEDS

Before performing the core functional connectivity analysis between entorhinal seeds and hippocampal voxels, we had to determine the entorhinal seeds, that is, the functional subregions of the entorhinal cortex (EC). We largely followed Maass, Berron et al. (2015) approach to assure comparability of results. The seeds were determined based on their functional connectivity with functionally and clinically relevant sources from the cortical item and contextual information processing streams, that are the perirhinal Area 35 and Area 36, the parahippocampal cortex and the retrosplenial cortex (see Nilssen et al., 2019).

The CONN toolbox (Whitfield-Gabrieli & Nieto-Castanon, 2012) was applied to perform a seed-to-voxel semipartial correlation analysis on the residual fMRI data between the retrosplenial, parahippocampal, Area 35 and Area 36 sources and the voxels within the segmented EC mask of each individual (see the description of the core functional connectivity analysis for the precise parameters). The resulting z-transformed correlation maps were then aligned for each participant to the group template T1 space and subjected to four one-sample T-tests (one for each source preference map) to reveal significant clusters of entorhinal connectivity preferences per source across all other entorhinal seeds, respectively. The functional subregions in the EC that we identify on group level generally overlap for the preferences towards the perirhinal cortex (Area 35 and Area 36) and towards the parahippocampal cortex with the findings by Maass, Berron et al. (2015). The exact procedure to determine the entorhinal seeds for further analysis is described in the main article.
